# Shared Fine-Scale Functional Organization in the Human Subcortex Revealed by Connectivity Hyperalignment

**DOI:** 10.64898/2026.09.29.755436

**Authors:** Jiaxin Cindy Tu, Ma Feilong, Yuqi Zhang, M Ida Gobbini, James V. Haxby

**Affiliations:** Department of Psychological and Brain Sciences, Dartmouth College; Department of Psychology, University of South Carolina; Department of Scientific Medicine and Surgery, University of Bologna

## Abstract

Subcortical structures are central to cognition, emotion, and social processing, and their dysfunction underlies a wide range of neurological and psychiatric conditions. Yet subcortical fine-scale functional organization has remained largely inaccessible to group-level fMRI analysis: nuclei are small and have low-SNR, and their functional topography varies idiosyncratically across individuals in ways that anatomical alignment cannot resolve. This idiosyncrasy carries real stakes. For example, deep brain stimulation targets with more variable functional connectivity across individuals show correspondingly less consistent clinical outcomes. Here we show that this seemingly idiosyncratic subcortical organization in fact reflects shared, fine-grained structure that can be recovered by modeling each individual’s connectivity in a common functional space (connectivity hyperalignment, CHA), rather than relying on anatomical correspondence. Applying CHA to fMRI data collected during naturalistic movie viewing, we found that aligning subcortical voxels by their cortical connectivity, rather than their anatomical location, sharply increased cross-subject correspondence in both stimulus-evoked responses and connectivity profiles. Critically, this alignment sharpened rather than blurred local structure: neighboring voxels became more functionally distinct from one another, and cortical connectivity targets became substantially more decodable across subjects. These results indicate that human subcortical nuclei possess a common fine-scale functional architecture that is masked by anatomical variability but recoverable at the individual level. This reframes a central methodological constraint in subcortical neuroimaging: rather than trading spatial resolution for signal quality via smoothing, group-level analyses can achieve voxel-level precision by aligning function rather than anatomy, opening a path to studying cortical-subcortical circuit organization in greater detail.

## Introduction

The human subcortex contains a diverse set of nuclei that are central to human cognitive, emotional, and social processing, and disruptions in their structure or function have been linked to a wide range of clinical conditions (Koshiyama et al., 2018; Okada et al., 2023). For example, the cortico-basal ganglia-thalamo-cortical loop is important for decision-making (Cruz et al., 2023; Tekin & Cummings, 2002), and the basal ganglia are heavily involved in the pathology of Parkinson’s disease (PD) (DeLong, 1990; DeLong & Wichmann, 2015; Graybiel, 2000; Obeso et al., 2000; Redgrave et al., 2010). Within the subcortical nuclei, there exists topographic organization of functional subdivisions that vary in their connectivity to cortical areas, as shown by early evidence from histology (Amunts et al., 2005; Kedo et al., 2018; Krauth et al., 2010; Morel et al., 1997; Yelnik et al., 2007), tract-tracing in animal studies (Freese & Amaral, 2006; Goldman-Rakic & Porrino, 1985; Haber, 2003; Middleton & Strick, 2002; Turner et al., 1980; Yeterian & Pandya, 1985), and non-invasive human neuroimaging (Behrens et al., 2003; Choi et al., 2012; Gordon et al., 2022; Hwang et al., 2017; Ji et al., 2019; Johansen-Berg et al., 2005; Jung et al., 2014; Keuken & Forstmann, 2015; O’Rawe & Leung, 2022; Pauli et al., 2018; Roy et al., 2009; Seitzman et al., 2020; Tian et al., 2020; Tyszka & Pauli, 2016). Beyond these discrete subdivisions, emerging evidence also reveals continuous connectivity gradients within the subcortex, with particularly detailed characterization in the thalamus (Oldham et al., 2025; Park et al., 2024; Sydnor et al., 2025; Wu et al., 2024). Converging evidence suggests that thalamocortical connectivity varies in its maturational timeline along a gradient that follows the sensorimotor-association axis of its cortical targets (Park et al., 2024; Sydnor et al., 2025). An analogous hierarchical organization extends across the broader subcortex, structuring both intrinsic neural timescales (Raut et al., 2020) and the heritability of functional gradients (Wu et al., 2024) in the striatum, thalamus, cerebellum, and hippocampus. Taken together, there is strong evidence that fine-scale functional organization pervades the human subcortex, manifesting both as discrete, nucleus-based subdivisions and as continuous connectivity gradients that traverse nuclear boundaries. However, like cortical organization, the subcortical functional topography possesses some idiosyncrasy across subjects, both in resting-state connectivity structure (Greene et al., 2020; Sylvester et al., 2020; Tian et al., 2020), and in task-evoked representational content (Sun et al., 2025).

This idiosyncrasy has real clinical consequences: deep brain stimulation targets with consistent functional connectivity (FC) across individuals, such as the thalamic motor integration zone used for essential tremor, produce reliably successful outcomes. On the other hand, targets with more idiosyncratic connectivity, such as the globus pallidus internus used for PD and dystonia, show far more variable clinical success (Greene et al., 2020). Consistent with this, in a cohort of PD patients treated with MRI-guided focused ultrasound, the distance between each patient’s lesion site and their own individually mapped thalamic functional target predicted treatment outcome (Ren et al., 2026). Modeling the shared structure underlying this fine-scale organization across individuals could therefore help to interpret the results from different patients.

Because of the small size, low signal-to-noise ratio (SNR), and close proximity of subcortical nuclei, visualizing and analyzing their fine-scale organization with MRI is challenging (Forstmann et al., 2017; Hollander et al., 2015). Spatial smoothing is often applied in fMRI analyses of the subcortex to compensate for low SNR and registration errors across individuals, which comes at the cost of disrupting the fine-scale structure within nuclei and mixing activity from multiple neighboring nuclei (Forstmann et al., 2017; Hollander et al., 2015). One promising approach for aligning fine-scale functional patterns in subcortical structures across brains is connectivity hyperalignment (CHA; Guntupalli et al., 2018; Haxby et al., 2020; Jiahui et al., 2023; Y. Zhang et al., 2026), which uses each voxel’s connectivity to cortical targets, rather than anatomical correspondence, to align patterns at this spatial scale. CHA projects individual connectivity data into a common model connectome (a group functional template) to model shared information embedded in idiosyncratic fine-scale multivoxel patterns. This procedure effectively aligns shared information embedded in fine-grained cortical functional topographies across individuals, facilitating group multivariate pattern analysis (MVPA) and predicting functional topography for an unseen individual (Guntupalli et al., 2018; Jiahui et al., 2023).

We hypothesize that CHA can similarly uncover shared structure in idiosyncratic subcortical topography, enabling group-level, individually predictive models of subcortical organization despite the region’s intrinsically lower SNR. Modeling this shared fine-scale structure would open new opportunities to study subcortical organization at a resolution previously inaccessible to group-level analysis. For example, group-level MVPA enabled by this shared structure could increase statistical power for delineating distinct input and output pathways to and from subcortical nuclei and their functional roles in the human brain, analogous to circuit-level insights obtained in animal studies (Ciocchi et al., 2010; Li et al., 2013; Orsini et al., 2011; Silberberg & Bolam, 2015). We test this hypothesis first in two smaller movie-viewing datasets (10-subject template, 13-subject test) and then replicate our key findings in an independent, higher-resolution 7T sample with substantially larger template and test sets (144 subjects for building a template and 34 subjects for validation testing).

## Results

### Connectivity hyperalignment (CHA) improves intersubject correlation of connectivity profiles and response time-series in subcortical regions

We applied connectivity hyperalignment (CHA) to project individual participants’ subcortical voxel spaces into a common connectome space (also known as the “group template”). We chose connectivity, rather than response profiles, as the alignment target for subcortical hyperalignment because subcortical representational geometry was far less consistent across subjects than cortico-subcortical connectivity geometry (**Supplementary Note 1**, **Supplementary Figures 1-3**). We further selected ridge-regression-based warping with cross-validated regularization (ridgeCV) over classic Procrustes alignment. Because connectivity geometry was only moderately reliable even within subjects across runs, a rigid transformation risked fitting run-specific noise rather than shared structure, and Procrustes hyperalignment failed to generalize to held-out runs where ridgeCV succeeded (**Supplementary Note 1**, **Supplementary Figure 4**).

The common connectome space was developed from data collected in 10 participants watching 50 minutes of the movie “The Grand Budapest Hotel” based on patterns of connectivity to 1210 cortical targets (ico8 resolution vertices on onavg cortical surfaces (Feilong et al., 2024)). Transformation matrices to the common model space were calculated in 13 other participants using patterns of connectivity while watching a different movie (**Figure 1**, details in **Materials and Methods**). To evaluate the effectiveness of CHA in aligning functional profiles, we examined leave-one-out intersubject correlation (ISC) in cortico-subcortical connectivity with the hyperaligned cortical targets (ico32 resolution vertices on onavg cortical surfaces, **Supplementary Note 2**), and ISC in subcortico-subcortical connectivity with contralateral subcortical targets (**Supplementary Note 3; Supplementary Figure 5**). Mean ISC of cortico-subcortical connectivity across all subcortical nuclei increased from 0.464 to 0.745 (Δz = 0.460, 95% CI [0.438, 0.482], p < 0.001; **Figure 2A; Supplementary Figure 6A**), when modeled with a linear mixed-effects model with random intercepts for subjects and nuclei. Similarly, subcortico-subcortical connectivity increased from 0.328 to 0.790 (Δz = 0.730, 95% CI [0.709, 0.752], p < 0.001; **Figure 2B; Supplementary Figure 6B**). Lastly, this effect was not limited to connectivity: ISC in response time series to the movie also increased from 0.02 to 0.09 (Δz = 0.070, 95% CI [0.066, 0.074], p < 0.001; **Figure 2C; Supplementary Figure 7**). To test whether this gain reflects denoising by ridge regression rather than alignment of shared functional structure, we compared CHA against a cross-run self-transform null that applies the same regularization without any cross-subject alignment. The null increased ISC relative to anatomical alignment, but by less than CHA, indicating that denoising explains only part of the gain (**Supplementary Note 4**).

**Figure 1.**
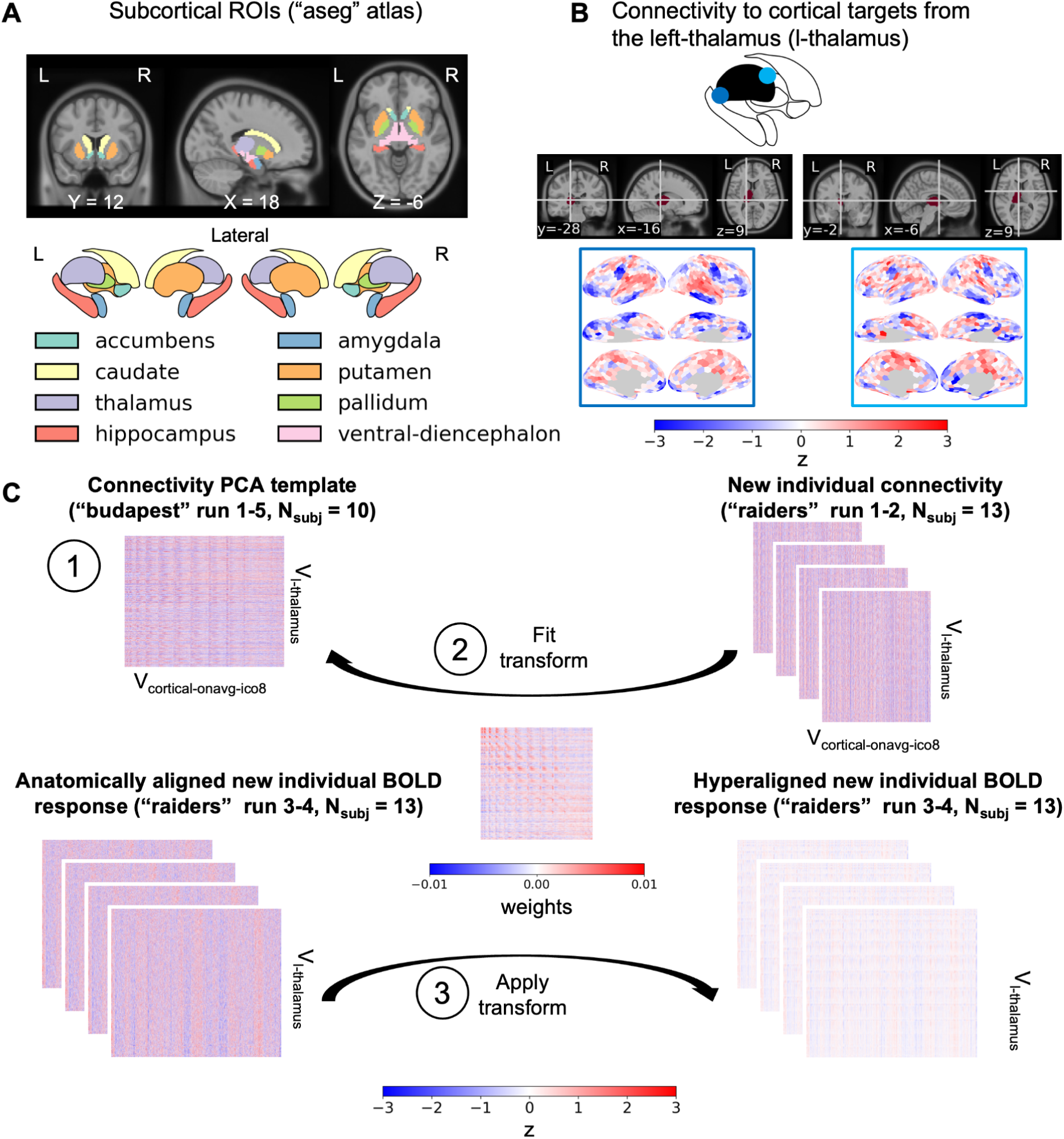
Connectivity hyperalignment of subcortical nuclei using functional connectivity (FC) to cortical targets (onavg-ico8 vertices). A) Visualization of 16 subcortical nuclei (8 each hemisphere) in coronal, sagittal and axial slices (left to right), and in a 2D vector outline plot below (ventral diencephalon omitted). B) Example connectivity patterns to onavg-ico8 cortical targets, shown for two different locations within the left thalamus. C) Hyperalignment procedure, illustrated using the left thalamus as an example. 1) Connectivities from voxels within the nuclei (V_l-thalamus_) to cortical onavg-ico8 targets (V_cortical-onavg-ico8_) were calculated in 10 participants (“budapest,” runs 1-5), and used to build a common model template. 2) The same connectivities (V_l-thalamus_ to V_cortical-onavg-ico8_) were calculated in an independent set of 13 participants watching the runs 1-2 of a different movie (“raiders”) and used to fit a transform aligning each individual’s left thalamus to the left thalamus template. 3) This transform was then applied to held-out data from the same 13 subjects collected while they watched runs 3-4 of the movie “raiders”). All validation testing was performed on these hyperaligned data. “budapest” refers to data collected while watching “The Grand Budapest Hotel” and “raiders” refers to data collected while watching “Raiders of the Lost Ark”. T = number of fMRI volumes/time points.

**Figure 2.**
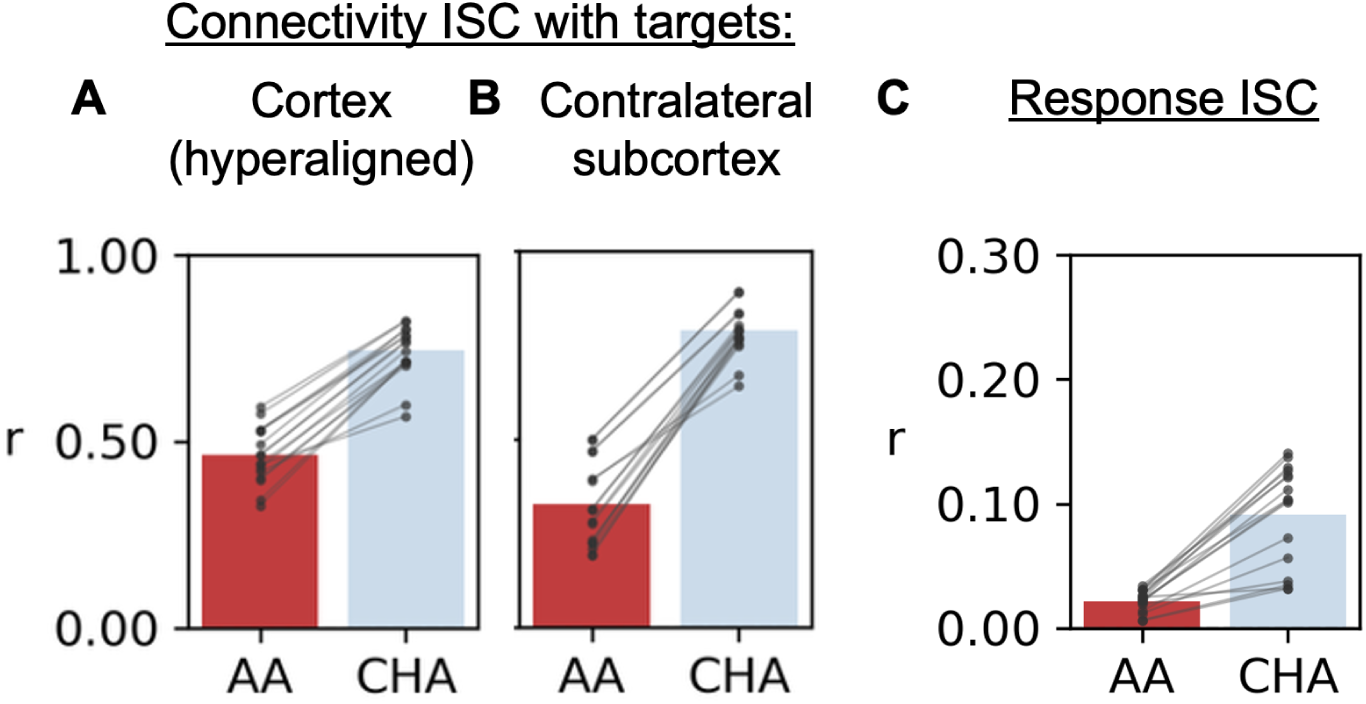
*Leave-one-out intersubject correlation (ISC) in subcortical functional profiles.* ISC in connectivity to A) hyperaligned onavg-ico32 cortical vertices, and B) contralateral subcortical voxels. C) ISC in movie responses. AA: anatomical alignment only. CHA: connectivity hyperalignment. Each data point represents statistics from a nucleus averaged across 13 participants.

### Connectivity hyperalignment in subcortical regions primarily affects the residual fine-grained responses and connectivity synchronization

To assess the effect of subcortical connectivity hyperalignment across spatial scales, we separated the movie responses into coarse-grained (nuclei means) and residual fine-grained (fine-grained multivoxel response patterns after subtracting the nucleus means) components (details in **Materials and Methods**). Here, coarse-grained connectivity is defined as correlation in time series between hyperaligned cortical onavg-ico8 vertices and the subcortical nuclei means (**Figure 3A)**, or between subcortical nucleus means and the contralateral subcortical nucleus means (**Figure 3B**). As expected from spatial pooling, anatomically aligned coarse-grained responses and connectivities were more synchronized across subjects than anatomically aligned fine-grained ones (compare **Figure 3A** to **3D**, **3B** to **3E**, **3C** to **3F**).

**Figure 3.**
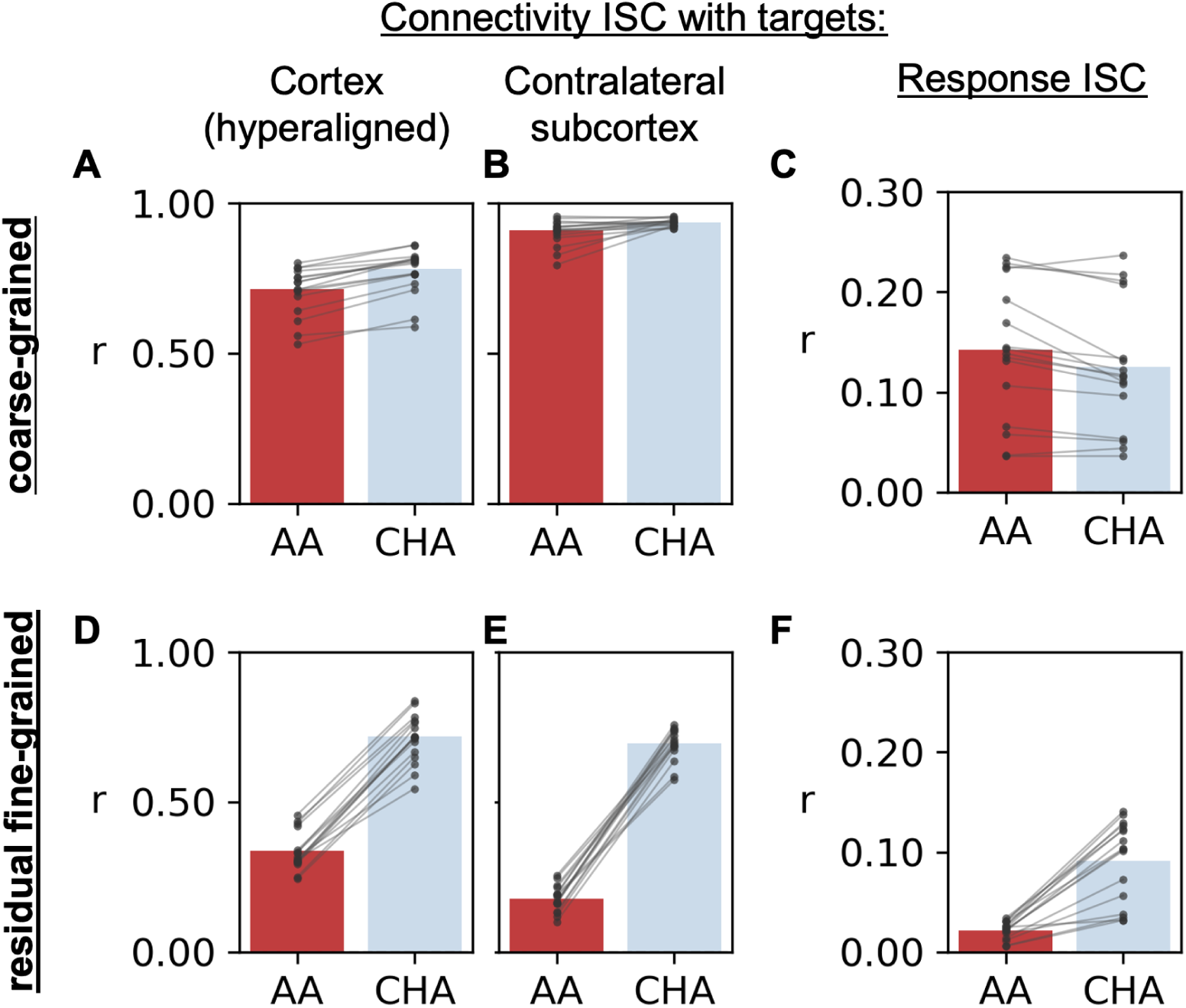
*Leave-one-out intersubject correlation (ISC) in coarse-grained and residual fine-grained subcortical functional profiles.* ISC in coarse-grained connectivity to A) hyperaligned cortical vertices (onavg-ico8), and B) contralateral subcortical nuclei means. C) ISC in coarse-grained movie responses. D-F) ISC in residual fine-grained connectivity and response after subtracting the nuclei mean response. AA: anatomical alignment only. CHA: connectivity hyperalignment. Each data point represents statistics from a nucleus averaged across 13 participants.

Overall, subcortical CHA modestly increased coarse-grained ISC in connectivity between subcortical nuclei means and hyperaligned cortical ico8 vertices from 0.713 to 0.780 (Δz = 0.152, 95% CI [0.130, 0.174], p < 0.001; **Figure 3A**); and between subcortical nuclei means to contralateral subcortical nuclei means from 0.909 to 0.936 (mean Δz = 0.186, 95% CI [0.132, 0.239], p < 0.001; **Figure 3B**). CHA slightly reduced coarse response ISC from 0.142 to 0.125 (Δz = −0.018, 95% CI [−0.023, −0.012], p < 0.001; **Figure 3C**). By contrast, CHA produced a large increase in residual fine-grained connectivity. ISC of residual fine-grained cortico-subcortical connectivity to hyperaligned cortical ico32 vertices increased from 0.337 to 0.720 (Δz = 0.557, 95% CI [0.533, 0.581], p < 0.001, **Figure 3D**). ISC of residual fine-grained subcortico-subcortical connectivity to contralateral subcortical voxels increased from 0.178 to 0.695 (Δz = 0.679, 95% CI [0.662, 0.696], p < 0.001, **Figure 3E**). ISC of residual fine-grained responses increased from 0.010 to 0.064 (Δz = 0.053, 95% CI [0.050, 0.057], p < 0.001, **Figure 3F**). Effects for individual subcortical nuclei are reported in **Supplementary Figures 10-12**. Since full fine-grained multivoxel patterns contain a mixture of coarse-grained and fine-grained information, subsequent analyses used residual fine-grained patterns to assess the spatial specificity of subcortical CHA.

### The spatial granularity of voxel functional profiles in the common model space

To assess whether the spatial granularity of subcortical functional profiles was preserved in the common model space, we computed the intersubject spatial point-spread function (PSF) for functional profiles using residual fine-grained responses and connectivities calculated with nucleus-mean subtracted time series. Briefly, instead of calculating ISC at the matching voxels across subjects, we calculated the ISC between a given voxel in one subject and neighboring voxels in other subjects at varying distances (binned to 2-8 mm in 1 mm intervals). Intuitively, if hyperalignment were simply smoothing or blurring signals across nearby voxels, this point-spread function would flatten (i.e., ISC would decay less steeply with distance) after CHA. In all subcortical nuclei after CHA, intersubject spatial PSFs in cortico-subcortical connectivities (**Figure 4A, 4B; Supplementary Figure 13**), or in subcortico-subcortical connectivities (**Figure 4C, 4D; Supplementary Figure 14**) were steeper (higher absolute values). Similarly, the intersubject spatial PSFs in response were also steeper (**Figure 4E, 4F; Supplementary Figure 15**). These results show that neighboring subcortical voxels have different functional profiles, and CHA models the commonality of these inter-voxel variations across brains.

**Figure 4.**
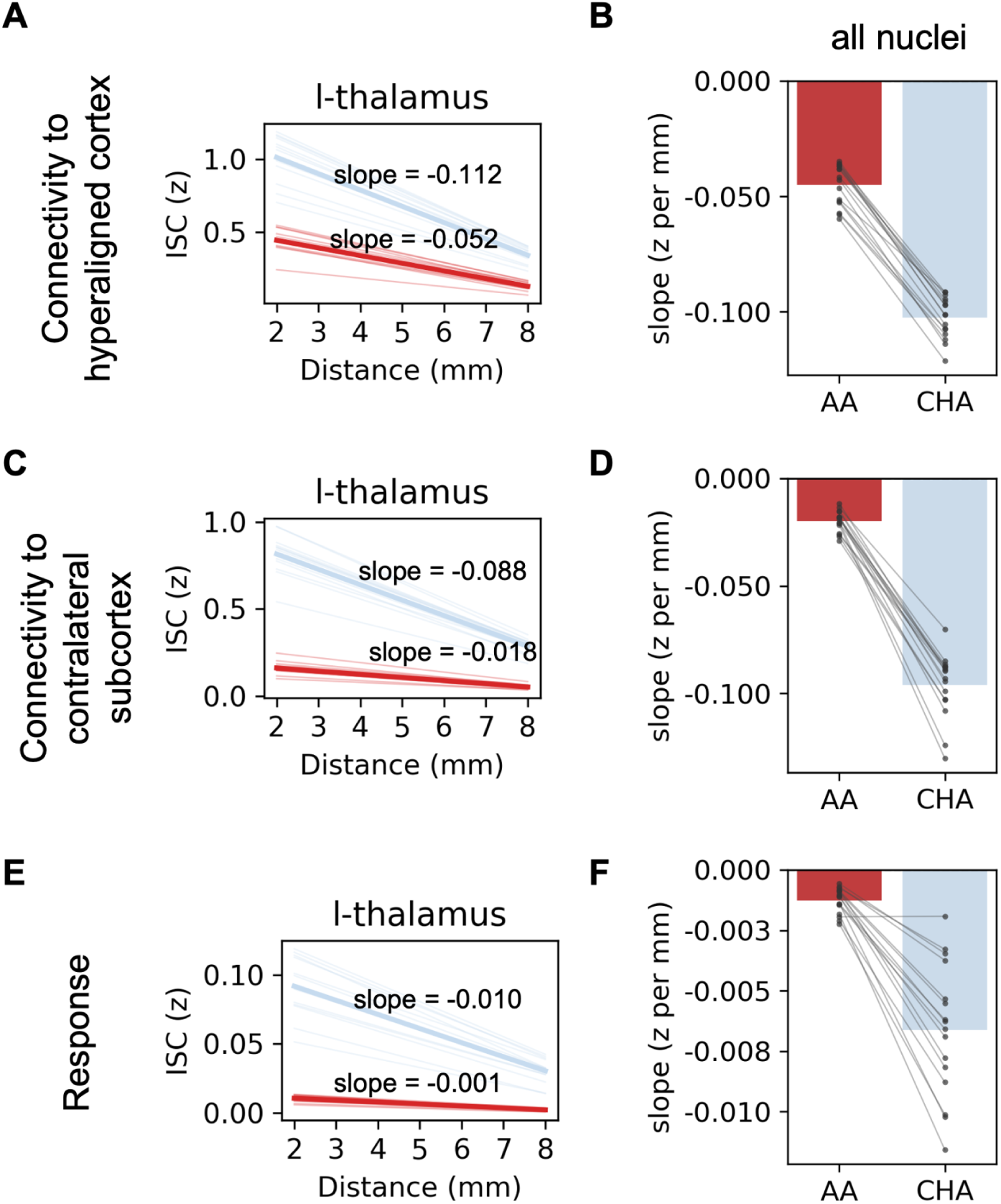
*Estimating the spatial granularity of shared representations in the common model space.* Intersubject spatial point spread function in ISC(z) of residual fine-grained A-B) connectivity to cortical targets, C-D) connectivity to contralateral subcortical targets, and E-F) responses. In A/C/E, left-thalamus (l-thalamus) is chosen as an example and each thin line is one subject, and the thick line is the average across subjects. B/D/F show the mean and standard error of slopes across 16 nuclei where each dot/line represents statistics from a nucleus averaged across 13 participants.

### Cross-subject prediction of connectivity targets based on hyperaligned patterns of connectivity within subcortical nuclei

Another test for spatial specificity of subcortical connectivity hyperalignment is whether projecting connectivity data into a CHA-derived common connectome space improves between-subject multivariate pattern classification of 1210 cortical connectivity targets (onavg ico8 vertices). For each subcortical nucleus, we predicted the cortical connectivity target based on the residual fine-grained multivoxel connectivity pattern (**Figure 5A**) using a nearest-neighbor classifier, where the predicted target was defined as the ico8 vertex whose average connectivity pattern across all other brains showed the lowest correlational distance with the example brain’s connectivity pattern (**Figure 5B-C**). Binary classification accuracy indexes whether the correlation distance is smaller for the correct target as compared to other single targets (pairwise classification, chance = 50%) and multiclass classification accuracy indexes whether the correlation distance is smaller for the correct target as compared to all 1209 other targets (chance < 0.1%). As seen in the example (**Supplementary Figure 16**), low correlational distances tend to come from ico8 vertices in the same cortical functional network (e.g. default mode network) as the target. Using a linear mixed-effects model, we found that relative to AA, CHA increased between-subject binary classification accuracy from 0.740 to 0.915 across targets (difference = 0.175, 95% CI [0.169,0.181], p < 0.001; **Figure 5D**), a level that is close to within-subject binary classification accuracy (≈ 0.938 across regions; **Supplementary Figure 17A**). Multiclass classification accuracy also increased from 0.017 to 0.070 across regions with CHA (difference = 0.053, 95% CI [0.047,0.059], p < 0.001; **Figure 5E**), close to within-subject multiclass connectivity target prediction (≈ 0.108 across regions; **Supplementary Figure 17B**).

**Figure 5.**
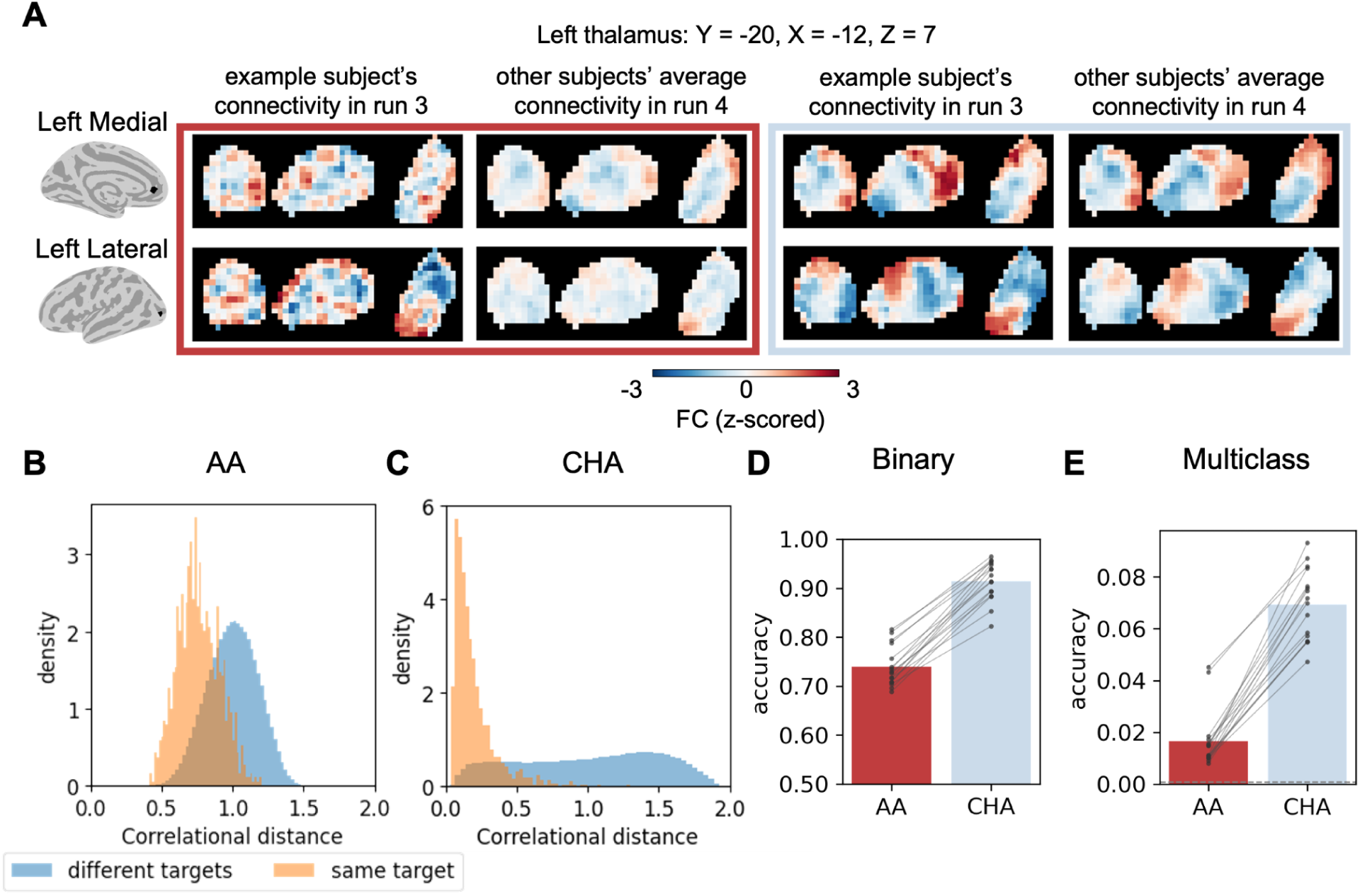
Cortical connectivity target (1210 onavg-ico8 cortical targets) prediction accuracies across subjects based on the residual fine-grained multivoxel pattern of functional connectivity (FC) in each subcortical nucleus after anatomical alignment (AA) and connectivity hyperalignment (CHA). A) Example connectivity patterns from the left thalamus to two cortical onavg-ico8 vertices located at left medial prefrontal cortex and left lateral occipital cortex in one example subject’s run 3 versus the average of all other subjects’ run 4. Coronal, sagittal and axial slices from left to right. Left (red) box: AA. Right (light blue) box: CHA. B) Correlational distance (1-Pearson’s r) between the AA connectivity profiles in example subject and all other subjects’ average (e.g. first and third columns in A) for between subject classification of cortical connectivity targets. Lower values indicate higher similarity. C) Same as B but for CHA connectivity profiles. D) Binary classification accuracy (chance = 0.5). E) Multiclass classification accuracy (chance = 1/1210 ≈ 0.0008, dashed line). Each data point represents statistics from a nucleus averaged across 13 participants. AA: anatomically aligned subcortical data and cortical data. CHA: connectivity hyperaligned subcortical data and cortical data.

### Hyperalignment weights favor spatially proximal voxels in subcortical nuclei

Because hyperalignment could in principle achieve its gains simply by pooling signal across a broad spatial neighborhood, we next asked how sharply the transformation weights are localized around each target voxel, and whether restricting hyperalignment to progressively smaller nuclei subdivisions degrades performance. Like cortical regions, subcortical nuclei can be further parcellated into smaller, functionally specialized nuclei. Tian et al. (2020) proposed a parcellation with four scales: 16 bilateral nuclei at scale 1, 32 at scale 2, 50 at scale 3, and 54 at scale 4 (**Figure 6A**). If the hyperalignment transformation pooled signals from only nearby voxels, then we would observe a large falloff in weights with distance, and restricting hyperalignment to the finer divisions (effectively setting the transformation weights outside the nuclei to zero) would not hurt the performance much.

**Figure 6.**
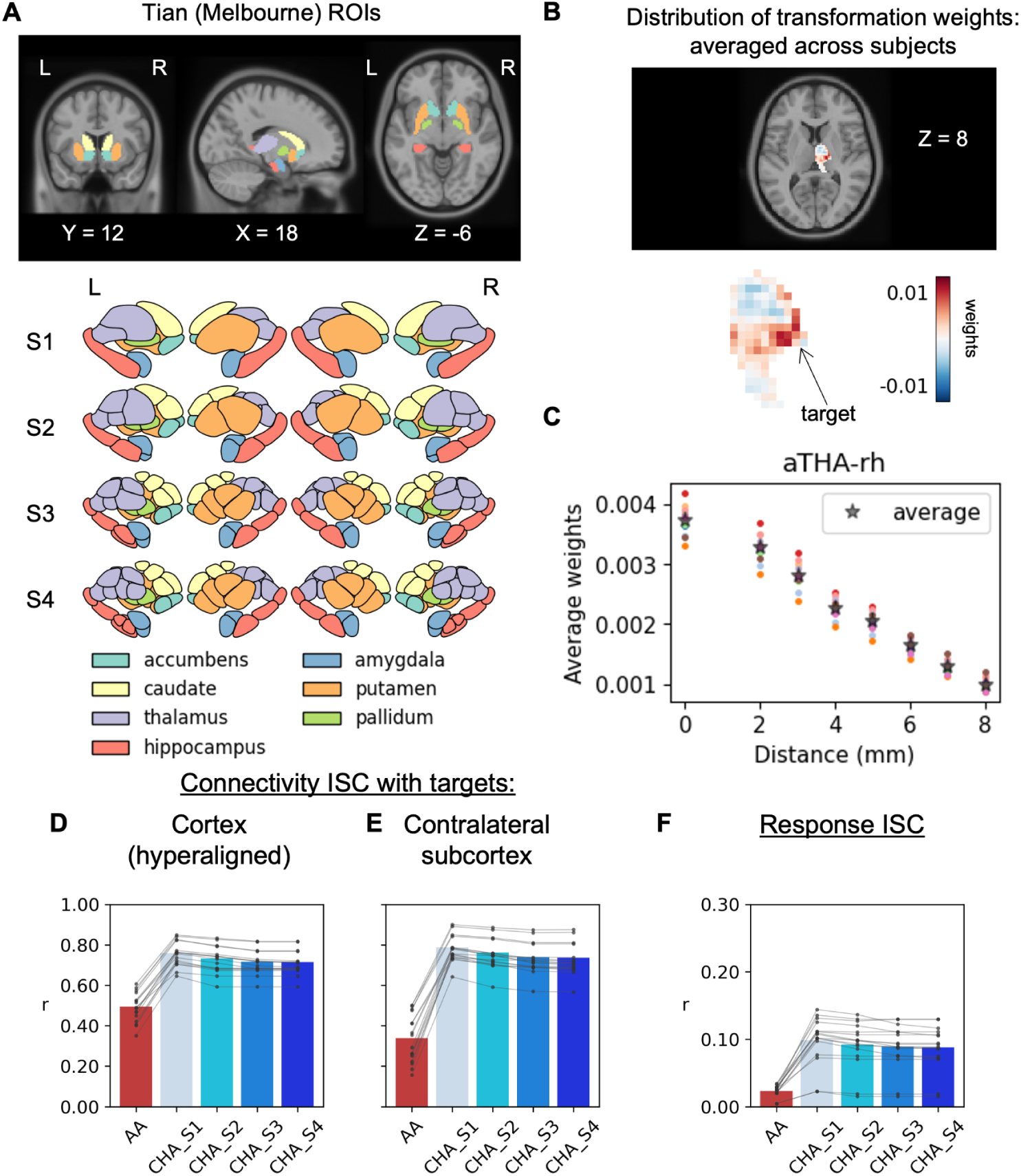
*Effect of the size of subcortical nuclei on hyperalignment performance.* A) The 7 core nuclei covered by the Tian atlas, shown in coronal, sagittal and axial views from left to right in the volumetric plot and from S1 to S4 in the schematic images. Note that S1 has 8 nuclei in each hemisphere because the thalamus is divided into anterior and posterior subnuclei. B) Distribution of transformation weights when hyperalignment transformation was calculated for the right anterior thalamus. C) Average weights by distance binned in 1 mm intervals. Dots represent individual participants and the star represents the average across individual participants. ISC in full fine-grained connectivity to D) hyperaligned cortical vertices (onavg-ico32), and E) contralateral subcortical voxels. F) ISC in full fine-grained movie responses. AA: anatomical alignment only. CHA: connectivity hyperalignment. Each data point represents one of 16 TianS1 nuclei (‘AMY-lh‘,’AMY-rh‘, ’CAU-lh‘, ’CAU-rh‘, ’GP-lh‘, ’GP-rh‘, ’HIP-lh‘, ’HIP-rh‘, ’NAc-lh‘, ’NAc-rh‘, ’PUT-lh‘, ’PUT-rh‘, ’aTHA-lh‘, ’aTHA-rh’,‘pTHA-lh‘, ’pTHA-rh’). AMY: amygdala, CAU: caudate, GP: (globus) pallidum, HIP: hippocampus, NAc: (Nucleus) accumbens, PUT: putamen, THA: thalamus. lh, left hemisphere; rh, right hemisphere

We first examined, for an example target voxel in the right anterior thalamus group template, how hyperalignment weights from all source voxels were distributed as a function of distance (averaged across participants; **Figure 6B**). Weights showed a negative relationship with Euclidean distance from the target voxel (**Figure 6C**), suggesting that spatially closer voxels were assigned larger weights. Across all 16 nuclei (defined at Tian S1), the slope in weights per mm ranged from −0.008 to −0.002.

Additionally, if we allow non-zero connectivity hyperalignment weights only within the smallest nucleus divisions (S4), we still observe a substantial increase with CHA compared to anatomical alignment in the ISC from 0.493 to 0.716 in cortico-subcortical connectivity (Δz = 0.358, 95% CI [0.341, 0.375], p < 0.001; **Figure 6D**), from 0.339 to 0.737 in subcortico-subcortical connectivity (Δz = 0.590, 95% CI [0.564, 0.616], p < 0.001; **Figure 6E**) and from 0.023 to 0.088 (Δz = 0.065, 95% CI [0.061, 0.068], p < 0.001; **Figure 6F**) in response time series. However, the benefit of CHA does decrease slightly with finer parcellations. For S1 relative to S4, ISC decreases from 0.759 to 0.716 in cortico-subcortical connectivity (Δz = −0.096, 95% CI [−0.105, −0.087], p < 0.001; **Figure 6D**); from 0.788 to 0.737 in subcortico-subcortical connectivity (Δz = −0.122, 95% CI [−0.134, −0.110], p < 0.001; **Figure 6E**), and from 0.098 to 0.088 (Δz = −0.011, 95% CI [−0.013, −0.009], p < 0.001) in response time series. Together, these results suggest that shared fine-scale topography remains detectable even within the smaller subdivisions defined by Tian S4.

### Application of hyperalignment to a large 7T dataset

We tested the generalizability of our approach with higher-resolution data collected in a larger sample, namely the Human Connectome Project (HCP) movie dataset collected with 7T scanning, 1.6 mm isotropic voxels, and 178 participants (Van Essen et al., 2013). In this analysis, the common connectome space template was built from 144 participants and evaluated on 34 participants (details in **Materials and Methods**). Despite the differences in acquisition resolution, task and nuclei definition (7T-version of the Tian atlas (Tian et al., 2020)), we replicated the increase in response and connectivity ISCs with CHA compared to AA, and the small decrease in ISC when CHA was restricted to finer spatial scales (from S1 to S4; **Figure 7**). The lower absolute ISC magnitude in HCP relative to raiders may be partly attributable to the discontinuous movie clip presentation in HCP versus the continuous movie presentation in raiders, and/or to differences in acquisition resolution (1.6 mm isotropic for HCP vs. 2.5 mm isotropic for raiders).

**Figure 7.**
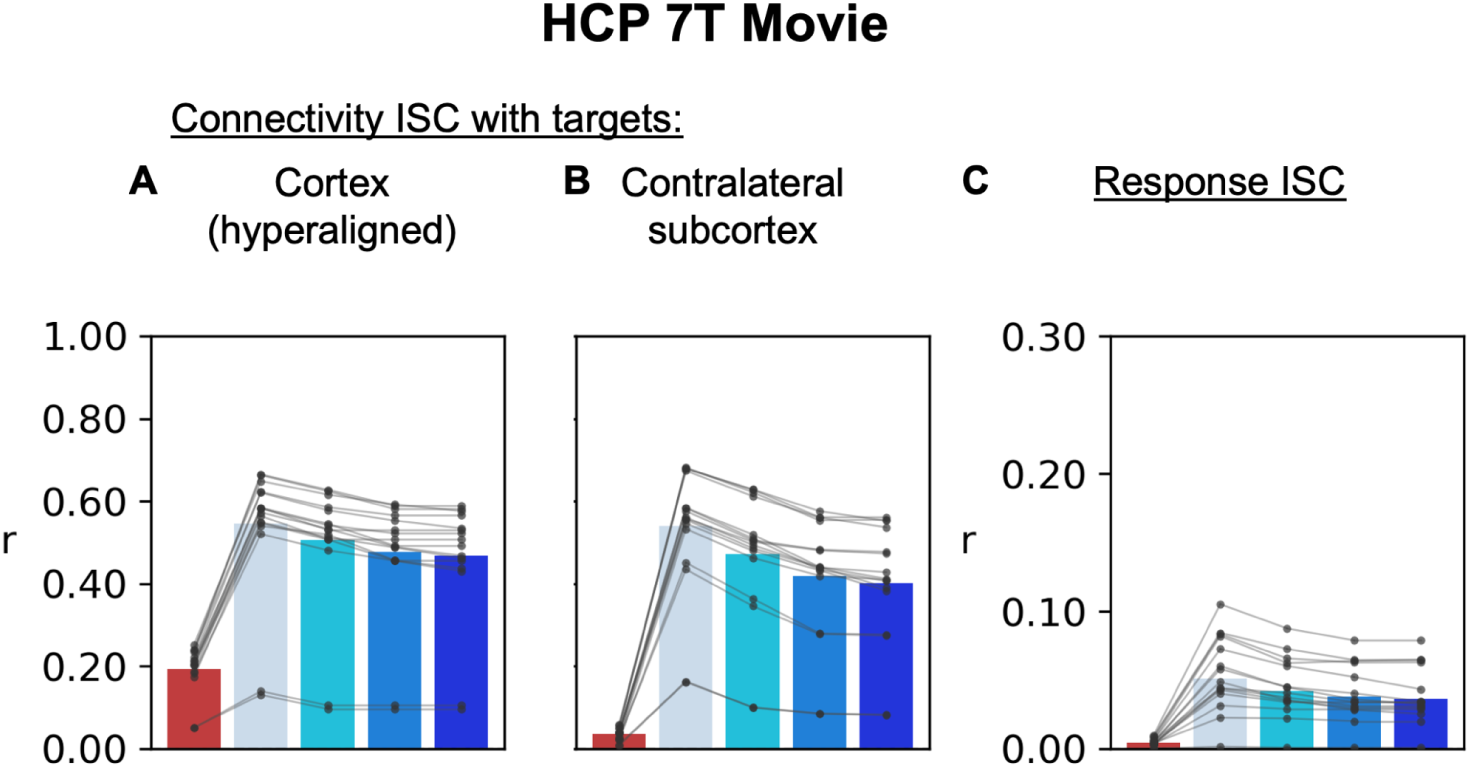
*Intersubject correlation (ISC) in response and connectivity for Tian (Melbourne) nuclei when connectivity hyperalignment was performed within nuclei at different spatial scales (S1 to S4).* AA: Anatomical alignment. CHA: Connectivity hyperalignment. Each line of data points represents one of the 16 Tian S1 nuclei.

## Discussion

The current study demonstrates that variable fine-grained connectivity patterns linking subcortical structures to cortical targets across individuals can be modeled using shared basis functions in a common model space. Hyperalignment to this space markedly increased cross-subject correlation of both functional connectivity and responses to movie stimuli, an effect largely driven by improved alignment of residual fine-grained structure rather than denoising. This finding suggests a need to re-evaluate current approaches to studying subcortical structures with fMRI: rather than relying on spatial smoothing, group-level analyses may be performed with greater precision at the voxel level by aligning fine-grained connectivity patterns across subjects.

### Connectivity hyperalignment as a versatile tool for aligning functional topography across subjects

Connectivity hyperalignment is a versatile approach that has several advantages over traditional response-based hyperalignment (Guntupalli et al., 2016; Haxby et al., 2011) for subcortical nuclei. First, representational geometry is relatively inconsistent in subcortical nuclei, making it challenging to build a common template that respects the central tendency of subcortical activation across subjects. On the other hand, there is evidence of widespread, converging, parallel connectivity between cortical regions and subcortical regions, with some clear topographic distributions (Alexander et al., 1986; Cambareri et al., 2025; Jallepalli & Dang, 2025). Second, while response hyperalignment requires a synchronized task such as a naturalistic movie-viewing paradigm, connectivity hyperalignment can be applied to movie-watching, rest and task data, and can therefore be used to re-analyze existing fMRI studies. Nevertheless, functional connectivity in movie-watching has previously been demonstrated to outperform resting-state functional connectivity in prediction of task activation and behavior (Finn & Bandettini, 2021; Gal et al., 2022), so it may still be desirable to use a rich naturalistic task when possible, for its ability to trigger a wide range of brain states. Third, connectivity across different scan runs provides repeated measurements for cross-validation-based hyperparameter tuning to optimize the hyperalignment transformation function without the need for repeated stimulus presentations.

### Connectivity hyperalignment captures the topographic organization of intrinsic connectivity

Decades of tract-tracing and neuroimaging studies have demonstrated that intrinsic connectivity within subcortical structures follows a topographic organization (Benarroch, 2015; Duvarci & Pare, 2014; Jallepalli & Dang, 2025; Ji et al., 2019; Øvsthus et al., 2024; Tian et al., 2020). Here, we observed that hyperaligned functional connectivity profiles were more similar across subjects at corresponding voxels while remaining distinct from those of neighboring voxels (**Figure 4**), suggesting that hyperalignment preserves fine-scale connectivity structure rather than blurring it as spatial smoothing would. Consistent with this, cross-subject decoding of cortical connectivity targets improved after hyperalignment: connectivity patterns linking subcortical voxels to different cortical targets became substantially more discriminable (**Figure 5**). Multiclass accuracy remained modest in absolute terms (≈7%), likely because cortical targets within the same functional network exhibit intrinsically similar connectivity patterns (**Supplementary Figure 16**). The single-nearest-target criterion for multiclass accuracy is a much stricter test than binary discrimination between two targets. Even within subjects across runs, the multiclass accuracy was only around 10%. The binary accuracy (≈ 92%) is near the ceiling of within-subject classification, which indicates that fine-scale connectivity differences are nonetheless reliably recoverable across subjects. These findings demonstrate the potential to identify subtle cortical-subcortical circuits in vivo using fMRI data aggregated across individuals.

### Limitations and future directions

While the current study establishes the presence of common fine-grained subcortical topography across subjects that can be recovered using connectivity hyperalignment, the extent to which this approach will advance our understanding of subcortical structures remains to be tested empirically, including the task-evoked maps. In addition, the performance of hyperalignment may improve with additional data both for constructing the group template and for estimating subject-specific transformations, as suggested by previous findings in cortical data (Feilong et al., 2023). Multi-echo EPI acquisition may further increase SNR in subcortical regions with short T2* such as the basal ganglia (Puckett et al., 2018).

Because comparisons were performed in the common model space rather than anatomical space, anatomical registration to a standard MNI template is in theory optional. When individual segmentations derived from high-resolution anatomical scans (e.g., 7T MRI) are available, connectivity hyperalignment could be performed directly in each subject’s native acquisition space, potentially minimizing registration error and partial-volume effects from resampling. As noted in a prior report, intersubject anatomical alignment can be particularly challenging for small nuclei or nuclei lacking clear anatomical landmarks (Kleckner et al., 2017). Alternatively, Tian and colleagues dilated group-level nuclei to create an uncertainty zone and applied support vector machine classification to derive individualized subcortical nuclei boundaries (Tian et al., 2020). Future work could examine whether improved nucleus definitions, or relaxing strict nucleus boundaries by allowing overlapping nuclei, may further enhance hyperalignment performance, analogous to the use of overlapping cortical searchlights in searchlight hyperalignment (Guntupalli et al., 2016, 2018; Haxby et al., 2020).

## Data availability

The Budapest dataset is available on OpenNeuro (https://doi.org/10.18112/openneuro.ds003017.v1.0.2) and the Raiders dataset will be made available on OpenNeuro.

## Code availability

The analysis code can be found at https://github.com/cindyhfls/Tu-2026-SubcortexCHA.

## Declaration of generative AI and AI-assisted technologies in the writing process

During the preparation of this work, the authors used Claude (Anthropic) to assist with editing and organizing manuscript text for clarity and readability, and to aid in the organization and documentation of analysis code (e.g., commenting, refactoring for clarity and efficiency, and reviewing code for potential errors). All scientific content, analyses, interpretations, and conclusions were developed by the authors. The authors reviewed and edited all AI-assisted material and take full responsibility for the content of the published article.

## Acknowledgements

The current study was funded by NIH R01MH127199.

## Materials and Methods

### Data acquisition

Two fMRI datasets were used for discovery analysis in which healthy adult human subjects watched naturalistic movies. fMRI data were collected while participants watched “The Grand Budapest Hotel” and “Raiders of the Lost Ark” at the Dartmouth Brain Imaging Center using a 3T Siemens Magnetom Prisma with a 32-channel head coil. For the functional runs, volumes were collected with 2.5 mm isotropic voxels and whole-brain coverage (TR = 1s, TE = 33 ms, flip angle = 59°, phase encoding direction anterior–posterior). Each volume comprised 52 axial slices collected in an interleaved fashion with gradient-echo echo-planar imaging. Each slice had a 96 × 96 matrix and an FOV of 240 × 240 mm. The imaging was accelerated using a simultaneous multislice (SMS) factor of 4 and no in-plane acceleration. The studies were approved by the Institutional Review Board of Dartmouth College. All participants gave written, informed consent, and were paid for their participation.

### The Budapest dataset (alias “budapest”)

The Budapest dataset consists of fMRI data from 25 participants (13 female, 24-31 years, mean age = 27.52 years) viewing the second half of the movie “The Grand Budapest Hotel” for ∼50 minutes, divided into 5 runs ranging from 9 to 13 minutes per run (Visconti di Oleggio Castello et al., 2020). Each run was padded with a 10 s fixation period both at the beginning and the end of the run. Here, we used 10 participants from this dataset to build the group template (details in “Connectivity hyperalignment (CHA) procedure”). All 10 participants (3 females, mean age = 28.1 years) had custom-fitted CaseForge headcases (https://caseforge.co) to minimize head motion in the scanner. Participant demographics and data quality are reported in a previous publication (Visconti di Oleggio Castello et al., 2020).

### The Raiders dataset (alias “raiders”)

The Raiders dataset (Feilong et al., 2023; Nastase, 2018) contains fMRI data from 23 participants (mean age ± SD: 27.3 ± 2.4 years; 12 females) while they watched the second half of the movie “Raiders of the Lost Ark”. Twenty of the 23 participants also participated in the Budapest dataset. The participants watched the first half of the movie outside the scanner, and watched the second half of the movie divided into 4 runs of 850, 860, 860, and 850s. The beginning 10 seconds of runs 2 to 4 overlapped with the last 10 seconds of the previous run, and were removed from analyses. Here, we selected the 13 participants who did not overlap with the 10 participants from the Budapest dataset, whose data were used to build the group template, to test the effect of connectivity hyperalignment.

### HCP 7T movie dataset (alias “hcp-7T”)

To test the robustness of our results, we also analyzed the HCP 7T movie fMRI data, which is part of the Human Connectome Project (Van Essen et al., 2013). Data were collected from 184 healthy young adults (aged 22–35) scanned at the Center for Magnetic Resonance Research at the University of Minnesota. Of these, 178 subjects had complete functional data for all four movie-watching runs and were included in our analyses (144 subjects for template building and 34 for testing). The HCP data were acquired using protocols approved by the Washington University institutional review board, and written informed consent was obtained from all subjects. Structural data were acquired using a customized 3 Tesla Siemens Connectom Skyra scanner with a standard Siemens 32-channel RF-receive head coil. At least one 3D T1w MPRAGE image and one 3D T2w SPACE image were collected at 0.7 mm isotropic resolution. The HCP fMRI data were acquired using a 7 Tesla Siemens Magnetom scanner with the Nova32 32-channel RF-receive head coil. Data were collected using a multiband gradient-echo EPI sequence (TR = 1000 ms, TE = 22.2 ms, flip angle = 45°, FOV = 208 × 208 mm, matrix = 130 × 130, spatial resolution = 1.6 mm isotropic voxels, 85 slices, multiband factor = 5, iPAT = 2, partial Fourier = 7/8, echo spacing = 0.64 ms, bandwidth = 1924 Hz/Px). The direction of phase encoding alternated between posterior-to-anterior (PA) and anterior-to-posterior (AP) across runs. During functional MRI acquisition, subjects passively viewed a series of audiovisual movie clips across four functional runs (2 runs each day across 2 days), each approximately 15 minutes in duration. Each run consisted of 4 or 5 clips ranging from 1:03 to 4:19 min:s, separated by 20-second rest periods. The first and third runs contained clips from independent films available under Creative Commons license; the second and fourth runs contained clips from Hollywood films. Audio was delivered via Sensimetrics earbuds, and movies were presented in full-screen mode (21.8° W × 15.7° H). Stimuli were presented using E-Prime.

### fMRI Data preprocessing

The Budapest and Raiders fMRI data were preprocessed with *fMRIPrep* version 20.2.7 (Esteban, Blair, et al., 2018; Esteban, Markiewicz, et al., 2018), which is based on *Nipype* version 1.7.0 (K. Gorgolewski et al., 2011; K. J. Gorgolewski et al., 2018). This includes standard procedures such as motion correction, surface reconstruction and anatomical registration to templates. Detailed descriptions from *fMRIPrep* boilerplate are provided in the **Supplementary Materials (Supplementary Note 5)** for reproducibility. Lastly, cortical data were resampled using intermediate files from fMRIPrep’s work directory with custom python scripts using “1step_pial_overlap” to the onavg-ico32 cortical surface template (Feilong, 2024; Feilong et al., 2024), which resamples directly from the original BOLD files onto the cortical surface with a single interpolation step by composing all the pertinent transformations (i.e. head-motion transform matrices, susceptibility distortion correction when available, and co-registrations to anatomical and output spaces). For subcortical volumetric data, a similar resampling method (“1step_linear_overlap”) was used to resample BOLD files onto the MNI152NLin2009cAsym space and a generic subcortical mask was applied to extract the nuclei (see section “Definition of subcortical mask and atlas”). For each functional run, linear regression was applied with nuisance regressors including 6 head motion parameters and their temporal derivatives, framewise displacement, 6 aCompCor components from cerebrospinal fluid and white matter, and polynomial trends up to the 2nd order. The estimated nuisance signal was then subtracted from the brain data, and the resulting residuals were z-scored across time independently at each vertex/voxel to normalize signal variance.

The HCP fMRI data were first downloaded as minimally preprocessed data (**Supplementary Note 6**), resampled to the onavg-ico32 cortical surface template (Feilong, 2024; Feilong et al., 2024), and additional nuisance regression was applied. For each fMRI run, nuisance regressors included 18 motion parameters derived from head motion correction, mean cerebrospinal fluid signal, mean white matter signal, and second-order Legendre polynomials to account for slow scanner drift. Movie viewing blocks within each run were extracted (starting 5 s after each stimulus onset and ending 5 s after stimulus offset) and concatenated into a single time series, and a single ordinary least squares regression was fitted across the concatenated segments simultaneously. The estimated nuisance signal was then subtracted from the brain data, and the resulting residuals were z-scored across time independently at each vertex/voxel per fMRI scan run to normalize signal variance.

### Definition of subcortical mask and atlas

For the 3T fMRI data from the Budapest and Raiders datasets, we first used a subcortical mask defined by the 16 nuclei using the population-level FreeSurfer “aseg” segmentation mask (https://github.com/templateflow/tpl-MNI152NLin2009cAsym/blob/master/tpl-MNI152NLin2009c Asym_res-02_seg-aseg_dseg.nii.gz), including bilateral accumbens, amygdala, caudate, putamen, thalamus, pallidum, hippocampus and ventral diencephalon (**Figure 1**). This covers 10,152 voxels in total (MNI152NLin2009cAsym - 2 mm resolution) and is referred to as the “aseg atlas”. We additionally repeated some analyses using an alternative segmentation of subcortical nuclei defined by Tian et al. (2020). This mask was created based on the probabilistic Harvard–Oxford Subcortical Structural Atlas (https://fsl.fmrib.ox.ac.uk/fsl/fslwiki/Atlases), with additional thresholding and enforcement of left-right symmetry, as described in detail in the original paper. Bilateral thalamus, caudate, putamen, accumbens, pallidum, hippocampus and amygdala were included in this atlas, with 8,562 voxels in total (MNI152NLin2009cAsym - 2 mm resolution). These subcortical nuclei were further parcellated based on their whole-brain functional connectivity gradients into smaller divisions. Here we used the nuclei defined at different spatial scales (scale I: 16 regions; scale II 32 regions; scale III: 50 regions; scale IV: 54 regions) downloaded from https://github.com/yetianmed/subcortex/blob/master/Group-Parcellation/. Unless otherwise noted, analyses in Figures 1–5 use the 16 nuclei defined by the aseg atlas; Figures 6–7 additionally use the Tian atlas parcellation at finer spatial scales.

For the 7T fMRI data (HCP), we used the 7T version of the Tian atlas, covering bilateral thalamus, caudate, putamen, accumbens, pallidum, hippocampus and amygdala, with 13974 voxels in total (scale I: 16 regions; scale II: 34 regions; scale III: 54 regions; scale IV: 62 regions, MNI152NLin6Asym - 1.6 mm resolution).

We used a subcortical visualization toolbox (Bryant, 2026) and nilearn (Nilearn contributors, n.d.) to visualize the individual nuclei divisions in brain slices and 2D vectors.

### Connectivity hyperalignment (CHA) procedure

A graphic illustration of this procedure can be found in **Figure 1**. The connectome for connectivity hyperalignment was based on functional connectivity with cortical targets for hyperalignment of both the cortex and subcortex. We used z-scored correlations between profiles from vertices/voxels and cortical vertices on onavg surface templates (Feilong et al., 2024) at ico-8 resolution (N = 1,210) as measures of functional connectivity. We used connectivity, rather than response profiles, as the basis for subcortical hyperalignment, because representational geometry was far less consistent across subjects in subcortical nuclei than has been reported in the cortex (Guntupalli et al., 2018), making response hyperalignment unsuitable (**Supplementary Note 1**).

To hyperalign subcortical data, group connectome templates were calculated for each nucleus based on data from the first 10 subjects watching ∼50 minutes of the movie “The Grand Budapest Hotel”, referred to here as “budapest” (Visconti di Oleggio Castello et al., 2020). Local connectome templates were computed for each subcortical nucleus, defined using the “aseg” (Fischl et al., 2002) and “Tian” atlases (Tian et al., 2020). Shared fine-grained connectivity information was extracted by taking the first *v* (*v =* number of voxels in the nucleus) principal components of horizontally stacked connectome matrices. This information was then rotated into the central tendency of voxel topography with a Procrustes transformation of vertically concatenated copies of the matrix into a vertical stack of the original connectome matrices (Feilong et al., 2023). Essentially, this template preserves shared information in fine-grained voxel-by-voxel differences in connectivity and resamples that information into the central tendency for its distribution over individual brains. Subsequently, individual transformation matrices were calculated using data from 13 different subjects watching a different movie, “Raiders of the Lost Ark”, referred to here as “raiders”. Transformation matrices were calculated using connectomes based on responses to the first two runs (runs 1 and 2) of “raiders”. The time-series responses to the last two runs (runs 3 and 4) were then resampled into the common model template space using the transformation matrices calculated based on the first two runs. Validation tests were performed on the hyperaligned subcortical data from these last two runs.

To calculate the intersubject correlation (ISC) in connectivity to hyperaligned cortical data, we additionally hyperaligned each cortical hemisphere with searchlight-based connectivity hyperalignment (Feilong et al., 2023; Guntupalli et al., 2018). Overlapping cortical searchlights (radius = 20 mm) were centered at onavg-ico32 vertices (N = 9,675 for left hemisphere and N = 9,666 for right hemisphere). A searchlight template was computed for each searchlight. Overlapping searchlight templates were aggregated using a weighted average to derive a whole-hemisphere template. Subsequently, a local transformation matrix was calculated for each individual’s searchlight connectome that resampled that searchlight’s vertices into the template. Searchlight transformation matrices were aggregated to derive a whole-hemisphere transformation using ridge regression. A detailed description of this procedure can be found in our previous publication (Feilong et al., 2023); minor modifications relative to that procedure (omission of distance weighting and of the bootstrap ensemble method, for reproducibility) are noted in **Supplementary Note 1**. Finally, the whole-hemisphere transformation matrices were applied to the BOLD response data in the last two runs of the movie “raiders” to generate the connectivity hyperaligned (CHA) data used for performance evaluations.

We calculated transformation matrices using ridge regression with cross-validated regularization (“ridgeCV”). The hyperparameter alpha was selected independently per participant and per nucleus for the “aseg” atlas, and as a single value across nuclei for the “Tian” atlas, using nested cross-validation, but the results from **Figure 6** were replicated using per-nucleus alpha selection (**Supplementary Figure 18; Supplementary Note 7**). Comparison of warp (ride-regression) hyperalignment Procrustes hyperalignment showed a clear advantage for ridge regression (**Supplementary Note 1)**.

Replication in the 7T HCP movie data largely follows the same steps used for the 3T “budapest” and “raiders” data. We computed the group connectome template from data collected in 144 participants watching the v1 of the movie task, and used this template to calculate transformations for 34 participants watching the v2 of the movie task. BOLD response data were truncated for each run to retain only the responses during movie presentation (starting 5 s after each movie clip onset and ending 5 s after clip offset). For each of the 34 test participants, a hyperalignment transformation matrix was calculated with the first two runs and applied to align the BOLD response data in the last two runs.

### Intersubject correlation in fMRI BOLD response to movies and dense connectivity

We used leave-one-out intersubject correlation (loo ISC) as a primary measure of hyperalignment performance, calculated as the correlation between a participant’s data and the mean of other participants. For ISC of connectivity to cortical targets, we used connectivity to all ico32 vertices (N = 19,341) instead of the coarser connectivity to ico8 vertices used for calculating hyperalignment. For ISC of connectivity to subcortical targets, we used connectivity to all subcortical voxels (N = 10,152 if using aseg atlas and N = 8,562 if using TianS4 atlas). For each subcortical nucleus we calculated ISCs between the flattened matrices of all cortical targets × all voxels. We also tested ISC of response profiles, using the flattened matrices of all time-points × all voxels. We Fisher-z-transformed ISCs before averaging across participants and inverse Fisher-z-transformed the mean. We performed tests of statistical significance using the Fisher-z transformed correlations (ISC(z)).

To control for the different levels of noise across subcortical structures, we also calculated a within-subject correlation (WSC) of connectivity. This can serve as a reference for the intersubject correlations. For this analysis, we calculated the correlation in connectivity between two test movie runs (“raiders” run 3 and run 4) where run 4 either comes from the same subject (within subject), or from another subject (across subject). In this case, we used the pairwise instead of loo ISC to facilitate a relatively fair comparison between within and across subject correlations. For each subject, the intersubject correlation was computed by averaging the Fisher-z-transformed pairwise correlations between that subject and all other subjects, and then transforming the result back to an r value.

### Separating response and connectivity into coarse- and residual fine-grained components

To further assess the extent to which functional profiles in subcortical nuclei at different spatial scales change after connectivity hyperalignment, we parsed responses into coarse-grained and residual fine-grained components. The coarse-grained response in each nucleus is defined as the mean fMRI BOLD response time-series; residual fine-grained responses are obtained by subtracting the nucleus mean from each voxel’s response. Similarly, coarse-grained connectivity is the connectivity between the nucleus mean response and other nucleus means (subcortical) or ico-8 vertices (cortical); the residual fine-grained functional connectivity is the connectivity between the residual fine-grained voxel response patterns in each nucleus and the fine-grained voxel response patterns of other nuclei or cortical vertices. While previous studies have alternatively defined coarse-grained connectivity as nucleus-to-vertex connectivity and fine-grained connectivity as voxel-to-voxel (Feilong et al., 2018), or coarse-grained connectivity as nucleus-to-voxel connectivity and fine-grained connectivity as vertex-to-vertex connectivity (Busch et al., 2024), we found the use of nucleus-to-voxel connectivity misleading because it mixed coarse-grained and fine-grained information, and was defined as “coarse” (Feilong et al., 2018) or “fine” (Busch et al., 2024) in different studies.

### Intersubject spatial point spread function calculation

To assess the spatial resolution of the effect of hyperaligning the data into the common model space, we calculated the intersubject spatial point spread function (PSF). Essentially, a loo ISC was calculated as the correlation of residual fine-grained connectivity or response as described above with the residual fine-grained connectivity or response of neighboring voxels in other subjects at distances ranging from 2 to 8 mm in 1 mm intervals (Euclidean distance in the MNI152NLin2009cAsym 2 mm resolution template). This analysis was conducted only for distances within nuclei. For each subject, we calculated the average response or connectivity patterns of all other subjects in a nucleus. ISCs for all pairs of voxels that belong to each distance bin were then Fisher Z-transformed and averaged. A linear regression was fitted to the mean ISC(z) values by distance so that a slope and intercept were obtained for each subject, and we estimated the mean and standard error of the slopes across subjects. We repeated the analysis with full fine-grained data in the Supplementary Materials. As a control, we also calculated the same spatial PSF in connectivity profiles for the same subject across runs.

### Predicting cortical connectivity targets based on fine-grained connectivity patterns within a single subcortical nucleus

For each subcortical nucleus, we asked whether its fine-grained multivoxel connectivity pattern could identify the correct cortical connectivity target among 1,210 onavg ico8 vertices. First, we calculated one connectivity pattern for each run using “raiders” run 3 and run 4. For each subject, we computed the average connectivity pattern in run 4 for each cortical vertex across all other subjects, excluding the current subject. We then compared the index subject’s connectivity pattern in run 3 for each cortical target to the leave-one-subject-out averages for other subjects, using correlational distance, yielding a 1,210 × 1,210 representational dissimilarity matrix (RDM). For each true cortical target, the vertex in the group average with the lowest correlational distance was then taken as the predicted target.

Binary classification accuracy was defined as the proportion of pairwise comparisons in which the correlation was higher for the same target than correlation with a different target, averaging across all target pairs (chance = 50%). Multiclass accuracy was the comparison of each target to all other targets, in which a correct classification was defined as the correlation with the correct target being higher than all individual correlations with the other 1,209 targets (chance < 0.1%). To estimate a performance ceiling, we repeated the analysis using connectivity patterns from run 4 of the same subject instead of the leave-one-subject-out averages (“within-subject classification”).

### Statistical analysis and software

Statistical analyses were performed in Python (v3.13.5) using the SciPy (v1.16.3) and statsmodels (v0.14.6) libraries. Linear mixed-effects models were fitted using restricted maximum likelihood (REML) when modeling the effect of connectivity hyperalignment. Subject and nuclei were modeled as crossed random effects (random intercepts), with hyperalignment condition (AA vs. CHA) as the fixed effect of interest. Model optimization was performed using the L-BFGS algorithm. If ISC or spatial slope in ISC is the predicted variable, we first applied a Fisher Z-transform to bring the Pearson’s r values to a near-normal distribution and model the effects in the z values. Significance was defined as p < 0.05 for the fixed effect. Because each reported model tests a distinct, pre-specified comparison (e.g., a specific alignment condition within a specific measure and dataset) rather than a family of comparisons over the same hypothesis, and because effect sizes were uniformly large relative to their confidence intervals, we did not apply correction for multiple comparisons across models.

## Notes

### Competing Interest Statement

The authors have declared no competing interest.

### Summary of Updates

clean up language, typo and minor adjustments

