## Supplemental Materials for "Shared Fine-Scale Functional Organization in the Human Subcortex Revealed by Connectivity Hyperalignment"

### Supplementary Materials

#### Supplementary Note 1: Justification for connectivity hyperalignment and choice of transformation and regularization

As noted in **Materials and Methods**, we selected ridgeCV over Procrustes hyperalignment as the transformation for subcortical CHA. Here we detail the comparisons motivating this choice, along with our rationale for using connectivity, rather than response profiles, as the alignment target.

In cortex, both response hyperalignment (RHA) and connectivity hyperalignment (CHA) are viable, and RHA sometimes outperforms CHA in the alignment of response profiles (Guntupalli et al., 2018), reflecting reasonably consistent representational geometry across subjects. For subcortical structures, however, we found representational geometry to be far less consistent across subjects (mean intersubject correlation of representational geometry matrices = 0.12, range 0.02-0.24; **Supplementary Figure 1**), making RHA an unsuitable choice for subcortical alignment. By contrast, connectivity geometry, i.e. the similarity structure among cortical targets' connectivity patterns within a given nucleus, was substantially more consistent across subjects (mean = 0.36, range 0.23-0.46 across nuclei; **Supplementary Figure 2**), indicating that connectivity carries more shared, alignable structure subcortically than response profiles do. We therefore used connectivity, rather than response profiles, as the basis for subcortical hyperalignment, and evaluated which transformation and regularization strategy best exploited this shared structure.

To determine how consistent this shared connectivity structure is across runs within the same subject, and not just across subjects, we also calculated within-subject reliability. For each subcortical nucleus, we correlated each subject's connectivity geometry in runs 1-2 against every subject's connectivity geometry in runs 3-4 (including the same subject), yielding a subject × subject matrix per nucleus in which the diagonal reflects within-subject reliability across runs, and the off-diagonal reflects between-subject consistency (**Supplementary Figure 3**). Average within-subject reliability ranged from 0.49 in left accumbens to 0.80 in right thalamus across the 16 nuclei, while average between-subject consistency was consistently lower, ranging from 0.22 in the left accumbens to 0.42 in the right thalamus. Because within-subject reliability is well below 1 for every nucleus, a transformation fit on runs 1 and 2 alone will partly capture noise specific to runs 1 and 2 rather than only the subject's stable, shared structure. Together, these results indicate that connectivity geometry, while more consistent across subjects than representational geometry, is still only moderately reliable within a subject across runs. Given this, we predicted that a rigid, unregularized transformation (Procrustes) would be more prone to fitting run-specific noise than a regularized, more flexible alternative (ridge regression), and would therefore generalize less well to held-out data.

Classic hyperalignment implements a rigid improper rotation via Procrustes transformation (Haxby et al., 2011). We compared Procrustes-based hyperalignment to ridge-regression-based "warp" hyperalignment (Feilong et al., 2023), which allows scaling and shearing, using either a fixed regularization parameter (alpha = 1e3, the default used for cortical hyperalignment) or nested cross-validation to select alpha per nuclei ("ridgeCV"). Consistent with our prediction, Procrustes hyperalignment increased ISC over anatomical alignment on training runs, across all three measures, but this improvement did not hold up on test runs (**Training runs**: connectivity to hyperaligned cortical vertices ISC: $\Delta z = 0.533$, 95% CI [0.516, 0.549], $p < 0.001$; connectivity to contralateral subcortical voxels ISC: $\Delta z = 0.156$, 95% CI [0.143, 0.170], $p < 0.001$; response ISC: $\Delta z = 0.022$, 95% CI [0.021, 0.023], $p < 0.001$; **Test runs**: connectivity to hyperaligned cortical vertices ISC: $\Delta z = 0.012$, 95% CI [0.005, 0.020], $p = 0.001$; connectivity to subcortical voxels ISC: $\Delta z = -0.027$, 95% CI [-0.038, -0.016], $p < 0.001$); response ISC: $\Delta z = -0.004$, 95% CI [-0.004, -0.003], $p < 0.001$; **Supplementary Figure 4**). Because Procrustes has fewer free parameters than ridge regression, this is not simply a matter of excess flexibility; rather, it suggests that Procrustes is fitting structure in runs 1-2 that does not generalize to different runs within the same subject. Both ridge (alpha = 1e3) and ridgeCV, by contrast, outperformed anatomical alignment on test runs across all three measures, indicating that the added flexibility of ridge regression (scaling and shearing, rather than rotation alone) generalizes better than Procrustes for connectivity hyperalignment.

We compared the performance of ridge regression with a fixed default regularization parameter (alpha = 1e3) to cross-validated ridge regression (ridgeCV) in which we estimate alpha with nested cross-validation. RidgeCV gave lower ISCs for connectivity to cortex on training runs but higher ISCs on test runs. **Training runs**: connectivity to hyperaligned cortical vertices ISC: fixed alpha versus ridgeCV-estimated alpha: $\Delta z = -0.104$, 95% CI [-0.116, -0.092], $p < 0.001$; **Test runs**: connectivity to hyperaligned cortical vertices ISC: $\Delta z = 0.119$, 95% CI [0.107, 0.131], $p < 0.001$. On the other hand, connectivity to subcortex and response ISCs were higher in both training runs and test runs for ridgeCV. **Training runs**: connectivity to contralateral subcortical voxels ISC: $\Delta z = 0.096$, 95% CI [0.081, 0.110], $p < 0.001$; response ISC: $\Delta z = 0.048$, 95% CI [0.043,0.052], $p < 0.001$; **Test runs**: connectivity to contralateral subcortical voxels ISC: $\Delta z = 0.261$, 95% CI [0.245, 0.277], $p < 0.001$; response ISC: $\Delta z = 0.049$, 95% CI [0.045,0.053], $p < 0.001$ (**Supplementary Figure 4**). Consequently, we report ridgeCV as our subcortical CHA implementation throughout the main paper.

Relative to our previous publication (Feilong et al., 2023), we made two minor modifications when aggregating searchlight transformation matrices: we did not use a distance weight, as early exploration found it negatively impacted hyperalignment performance, and we did not implement the bootstrap-based "ensemble" method, as it yielded little performance improvement over a single fit at much higher computational cost.

#### Supplementary Note 2: Using hyperaligned cortical data versus anatomically aligned cortical data for intersubject correlations (ISC) in cortico-subcortical connectivity

We presented the results of ISC in cortico-subcortical connectivity with hyperaligned cortical data, but the results are qualitatively replicated with anatomically aligned cortical targets.

#### Supplementary Note 3: Rationale for using contralateral nuclei when calculating intersubject correlations (ISC) in subcortico-subcortical connectivity

Because the subcortical nuclei are small, within-nucleus functional connectivity (FC) estimates may be particularly sensitive to spatial smoothness, including that introduced by resampling to the MNI152NLin2009cAsym template at 2-mm resolution. Spatial smoothness and head motion can inflate correlations between nearby voxels, potentially increasing apparent intersubject correlation (ISC) of subcortical connectivity profiles. To assess the contribution of local connections, we compared ISC using all subcortical voxels as connectivity targets against three exclusion conditions for each of the 16 nuclei: (1) excluding targets within the seed nucleus; (2) excluding all ipsilateral targets, retaining only contralateral subcortical voxels; and (3) retaining only contralateral targets outside the homologous nucleus. Excluding the seed nucleus substantially reduced ISC, whereas excluding all ipsilateral targets yielded similar ISC values. Further excluding the contralateral homologous nucleus reduced ISC (**Supplementary Figure 5**). We therefore used all contralateral subcortical voxels as targets in subsequent subcortico-subcortical connectivity analyses. This approach excluded local connections and reduced computational demands while ensuring identical target counts across seed nuclei within each hemisphere and approximately equal target counts across hemispheres. By contrast, excluding only the seed nucleus would yield target counts that varied with nucleus size. Similar exclusions of local connections have been used to mitigate motion-related inflation of short-range FC (Power et al., 2011).

#### Supplementary Note 4:Is the ISC gain from CHA explained by denoising?

CHA increased ISC of connectivity, but this gain is hard to interpret without a reference for how high ISC could possibly be given measurement noise, and because the ridge regression used in CHA could raise ISC simply by denoising the data rather than by aligning shared functional structure. To address both issues, we first estimated a noise ceiling using the within-subject correlation (WSC) in connectivity between the two test runs (run 3 and run 4), where the run 4 connectivity came either from the same subject (WSC) or from a different subject (ISC). For a fair comparison, we used pairwise rather than leave-one-out ISC and a single run rather than both, so these values are lower than those reported in the main text. As expected, nuclei with higher WSC tended to have higher ISC (**Supplementary Figure 8**). However, WSC also increased after CHA, possibly because ridge regression favors more reliable voxels when mapping to the common space, which suggests that denoising could account for part of the ISC gain. To test this, we built a “cross-run self-transform” null in which the transformation was estimated between runs 1 and 2 of the same subjects, applying the same regularized mapping without any cross-subject alignment (**Supplementary Figure 9A**). This null increased ISC relative to anatomical alignment, indicating that denoising contributes, but the increase from real hyperalignment was larger (**Supplementary Figure 9B, left**). The gain also did not depend on building a template: transforming each subject directly into the reference subject’s space still raised ISC above both anatomical alignment and the cross-run null (**Supplementary Figure 9B, right**). Together, these results indicate that denoising explains part, but not all, of the ISC gain from CHA, and that the remainder reflects recovery of a shared, voxel-specific correspondence in fine-scale topography across subjects rather than a general reduction in noise.

#### Supplementary Note 5: fMRIPrep data processing

Anatomical data preprocessing: One T1-weighted (T1w) image was corrected for intensity non-uniformity (INU) with `N4BiasFieldCorrection`(Tustison et al., 2010), distributed with ANTs 2.3.3 (Avants et al., 2008), and used as T1w-reference throughout the workflow. The T1w-reference was then skull-stripped with a *Nipype* implementation of the `antsBrainExtraction.sh` workflow (from ANTs), using OASIS30ANTs as target template. Brain tissue segmentation of cerebrospinal fluid (CSF), white-matter (WM) and gray-matter (GM) was performed on the brain-extracted T1w using `fast` [FSL 5.0.9,(Zhang et al., 2001)].
Brain surfaces were reconstructed using `recon-all` [FreeSurfer 6.0.1,(Dale et al., 1999)], and the brain mask estimated previously was refined with a custom variation of the method to reconcile ANTs-derived and FreeSurfer-derived segmentations of the cortical gray-matter of Mindboggle (Klein et al., 2017). Volume-based spatial normalization to one standard space (MNI152NLin2009cAsym) was performed through nonlinear registration with `antsRegistration` (ANTs 2.3.3), using brain-extracted versions of both T1w reference and the T1w template. The following template was selected for spatial normalization:
*ICBM 152 Nonlinear Asymmetrical template version 2009c* [(Fonov et al., 2009); TemplateFlow ID: MNI152NLin2009cAsym],

Functional data preprocessing: For each of the BOLD runs found per subject (across all tasks and sessions), the following preprocessing was performed. First, a reference volume and its skull-stripped version were generated using a custom methodology of *fMRIPrep*. A B0-nonuniformity map (or *fieldmap*) was estimated based on a phase-difference map calculated with a dual-echo GRE (gradient-recall echo) sequence, processed with a custom workflow of *SDCFlows* inspired by the [`epidewarp.fsl` script](http://www.nmr.mgh.harvard.edu/~greve/fbirn/b0/epidewarp.fsl) and further improvements in HCP Pipelines (Glasser et al., 2013). The *fieldmap* was then co-registered to the target EPI (echo-planar imaging) reference run and converted to a displacements field map (amenable to registration tools such as ANTs) with FSL's `fugue` and other *SDCflows* tools. Based on the estimated susceptibility distortion, a corrected EPI (echo-planar imaging) reference was calculated for a more accurate co-registration with the anatomical reference. The BOLD reference was then co-registered to the T1w reference using `bbregister` (FreeSurfer) which implements boundary-based registration (Greve & Fischl, 2009). Co-registration was configured with six degrees of freedom. Head-motion parameters with respect to the BOLD reference (transformation matrices, and six corresponding rotation and translation parameters) are estimated before any spatiotemporal filtering using `mcflirt` [FSL 5.0.9 (Jenkinson et al., 2002)]. The BOLD time-series (including slice-timing correction when applied) were resampled onto their original, native space by applying a single, composite transform to correct for head-motion and susceptibility distortions. These resampled BOLD time-series will be referred to as *preprocessed BOLD in original space*, or just *preprocessed BOLD*. First, a reference volume and its skull-stripped version were generated using a custom methodology of *fMRIPrep*. Several confounding time-series were calculated based on the *preprocessed BOLD*: framewise displacement (FD), DVARS and three region-wise global signals. FD was computed following Power (absolute sum of relative motions, (Power et al., 2014)). FD and DVARS are calculated for each functional run, both using their implementations in *Nipype* [following the definitions by (Power et al., 2014)]. The three global signals are extracted within the CSF, the WM, and the whole-brain masks. Additionally, a set of physiological regressors were extracted to allow for component-based noise correction [*CompCor*, (Behzadi et al., 2007)]. Principal components are estimated after high-pass filtering the *preprocessed BOLD* time-series (using a discrete cosine filter with 128s cut-off) for the two *CompCor* variants: temporal (tCompCor) and anatomical (aCompCor). tCompCor components are then calculated from the top 2% variable voxels within the brain mask. For aCompCor, three probabilistic masks (CSF, WM and combined CSF+WM) are generated in anatomical space. The implementation differs from that of Behzadi et al. in that instead of eroding the masks by 2 pixels on BOLD space, the aCompCor masks are subtracted a mask of pixels that likely contain a volume fraction of GM. This mask is obtained by dilating a GM mask extracted from the FreeSurfer's *aseg* segmentation, and it ensures components are not extracted from voxels containing a minimal fraction of GM. Finally, these masks are resampled into BOLD space and binarized by thresholding at 0.99 (as in the original implementation). Components are also calculated separately within the WM and CSF masks. For each CompCor decomposition, the *k* components with the largest singular values are retained, such that the retained components' time series are sufficient to explain 50 percent of variance across the nuisance mask (CSF, WM, combined, or temporal). The head-motion estimates calculated in the correction step were also placed within the corresponding confounds file.

#### Supplementary Note 6: HCP minimal preprocessing

Similar to fMRIPrep data processing, the HCP minimal preprocessing (Glasser et al., 2013) involves passing the distortion- and bias-corrected T1w image to FreeSurfer and running “recon-all” to generate brain surfaces. The T2w image was is used to assist pial surface delineation by removing dura and blood vessels. Next, all anatomical surface files are converted from the subject's native brain space into a standard reference space (called MNI) using a nonlinear transformation. Functional data in this space were subsequently mapped to onavg-ico32 surfaces.

##### Supplementary Note 7: pooled versus separate alpha selection per nucleus

When using a pooled alpha selection, where a single alpha for ridge regularization was chosen for all nuclei, the alpha was consistently chosen to be 10000 regardless of participants and atlases used (“aseg” or “Tian” at S1/S2/S3/S4). However, when alpha was selected separately for each nucleus, the finer-scale atlases with smaller nuclei (e.g., S4) had more alpha chosen to be 1000. ISC in connectivity hyperaligned data using pooled alpha is consistently higher than that using separate alpha, but the trend of decreasing ISC from S1 to S4 still holds with separate alpha selection. (**Supplementary Figure 18**).

### Supplementary Figures

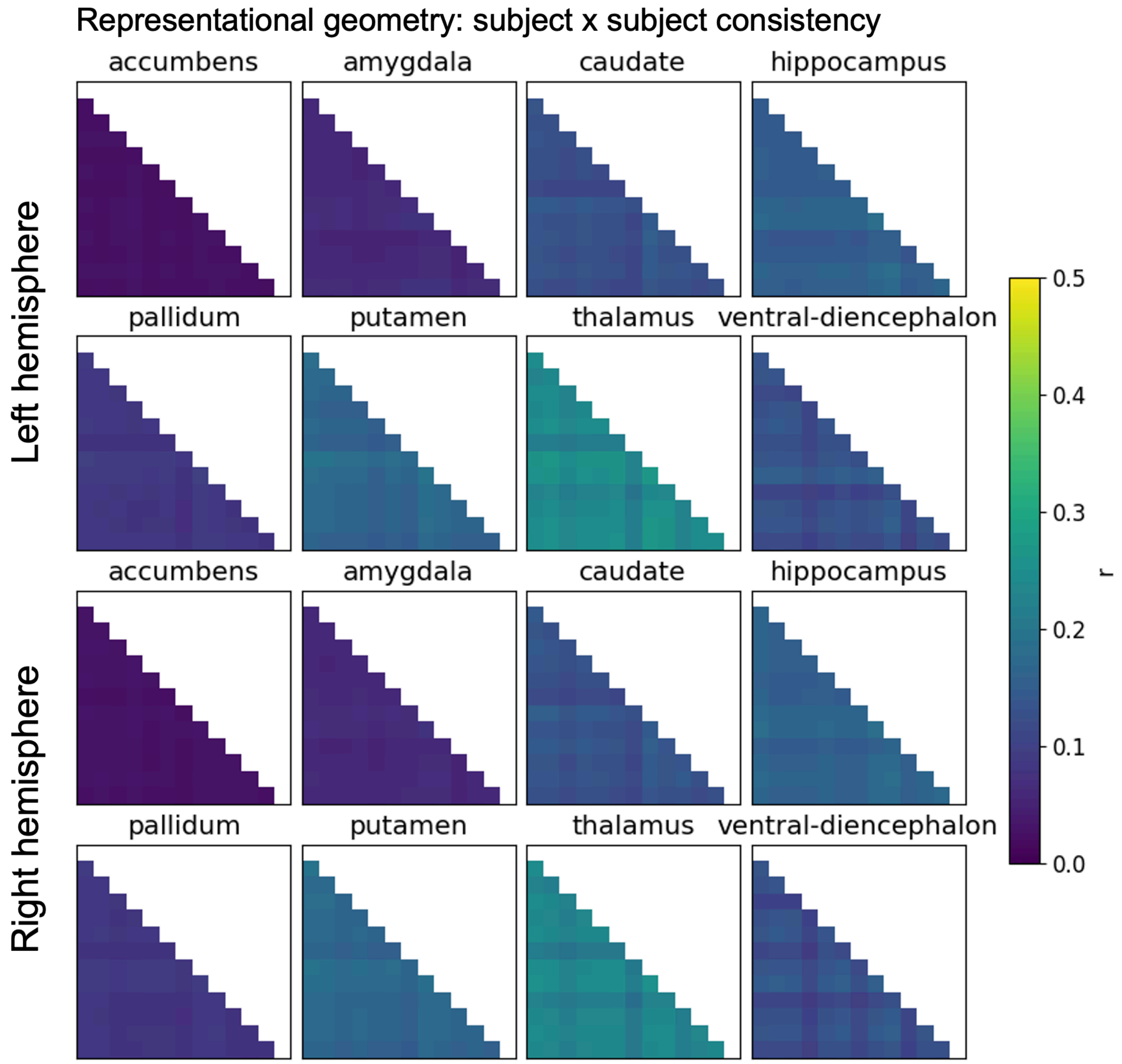


**Supplementary Figure 1.** *Representational geometry consistency in subcortical nuclei.*

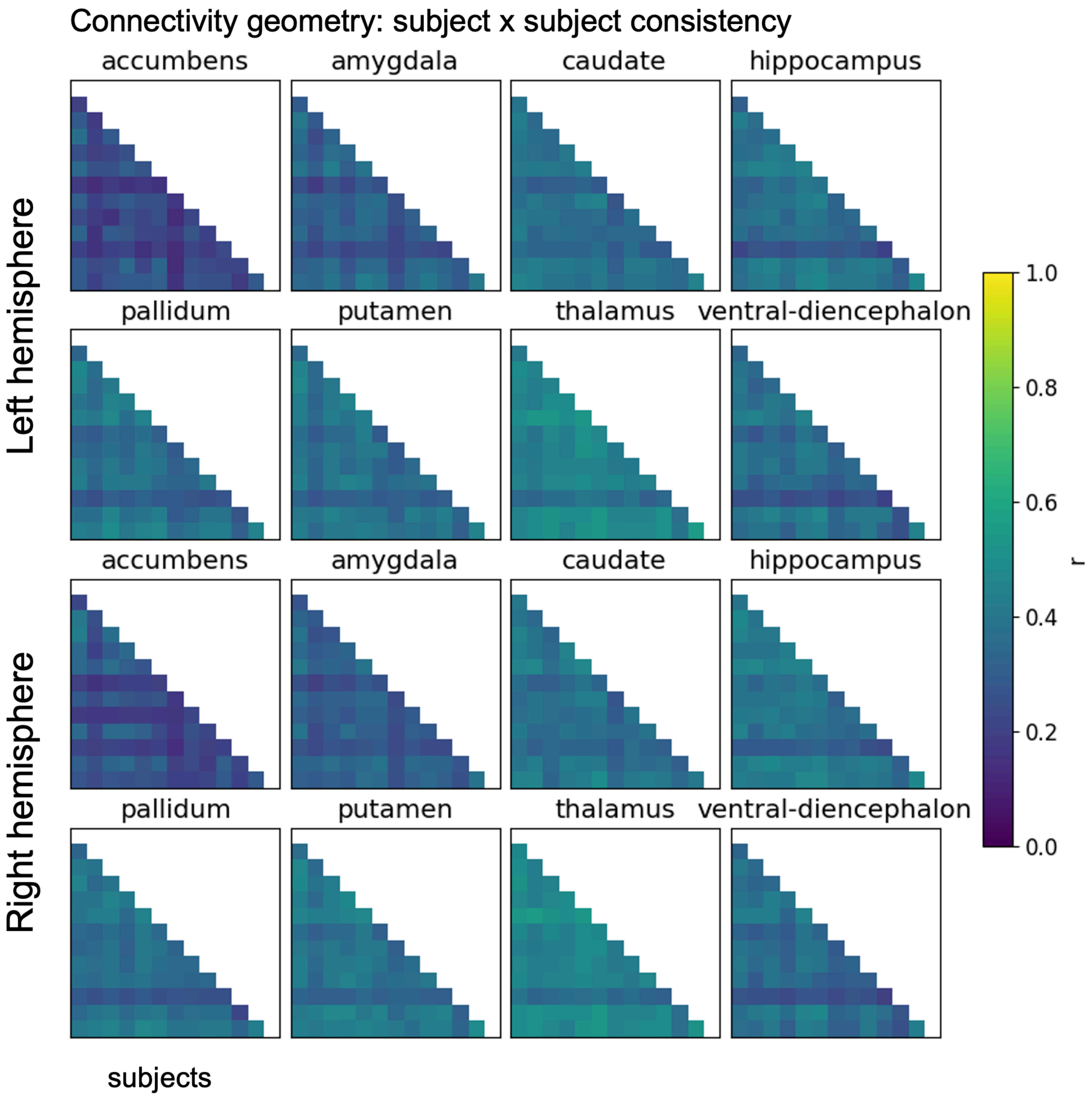


**Supplementary Figure 2.** *Connectivity (to cortical onavg-ico8) geometry consistency in subcortical nuclei.*

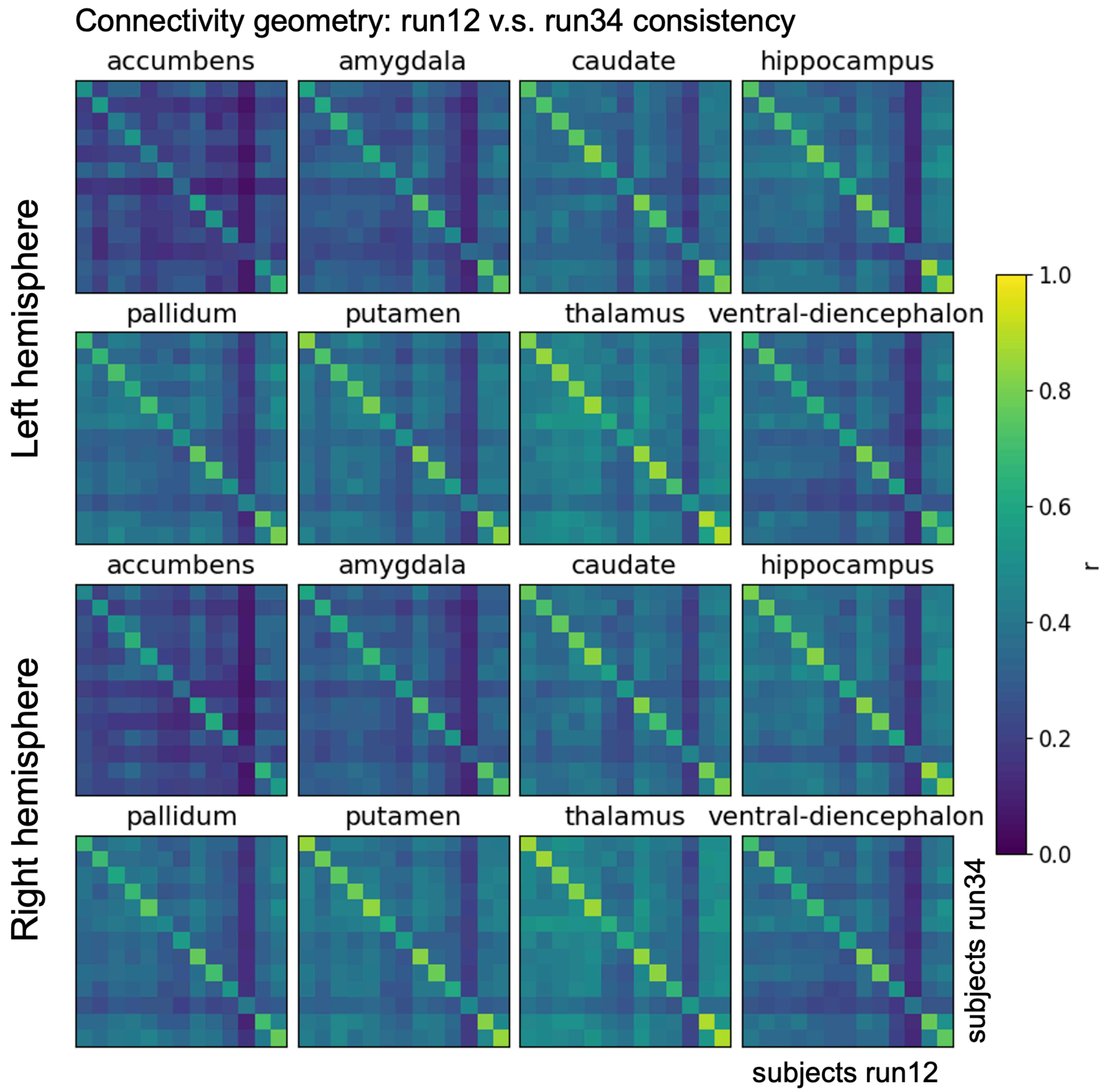


**Supplementary Figure 3.** *Connectivity (to cortical onavg-ico8) geometry consistency in subcortical nuclei between runs 1 and 2 of subjects to runs 3 and 4 of subjects.*

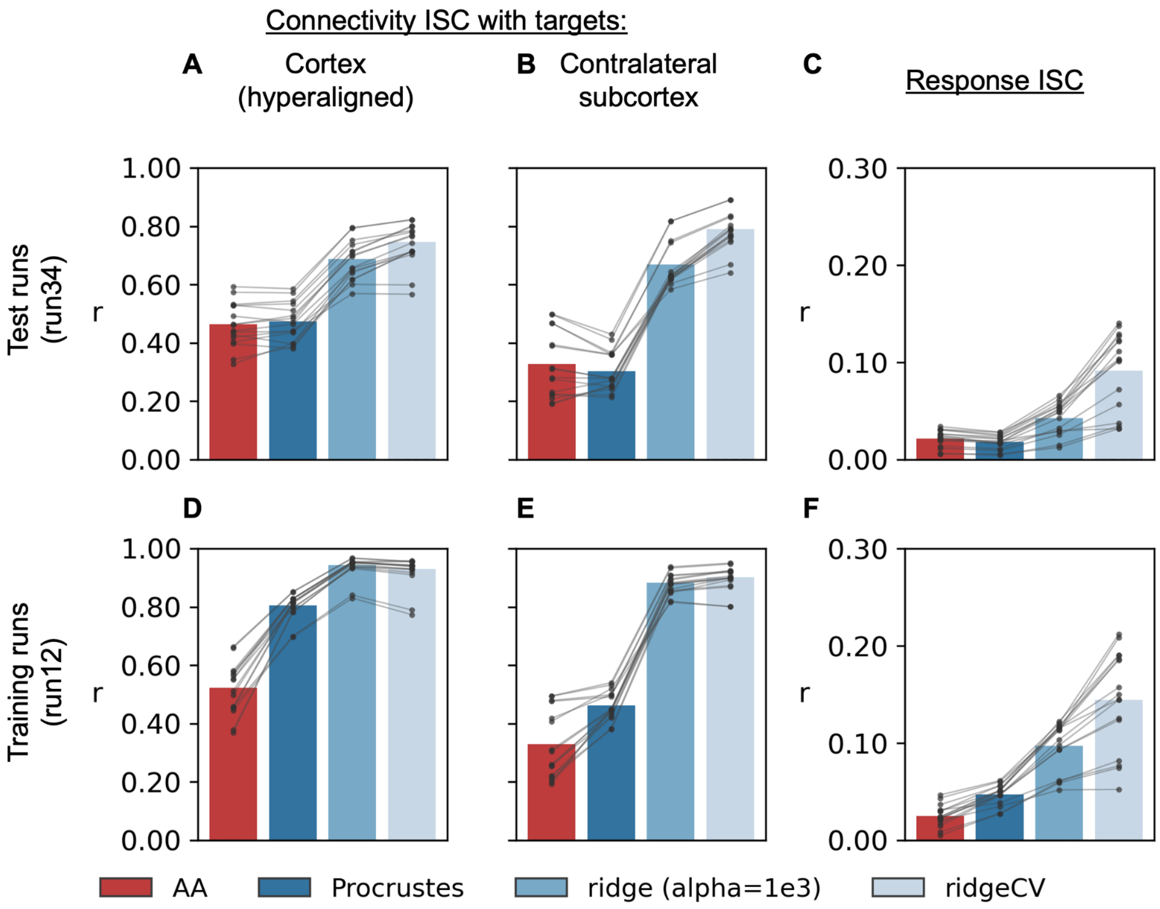


**Supplementary Figure 4***. Intersubject correlation (ISC) in connectivity and response profiles to all subcortical voxels and all hyperaligned cortical onavg-ico32 vertices.* AA: Anatomical alignment only. CV: cross-validation. Each data point represents one of 16 nuclei.

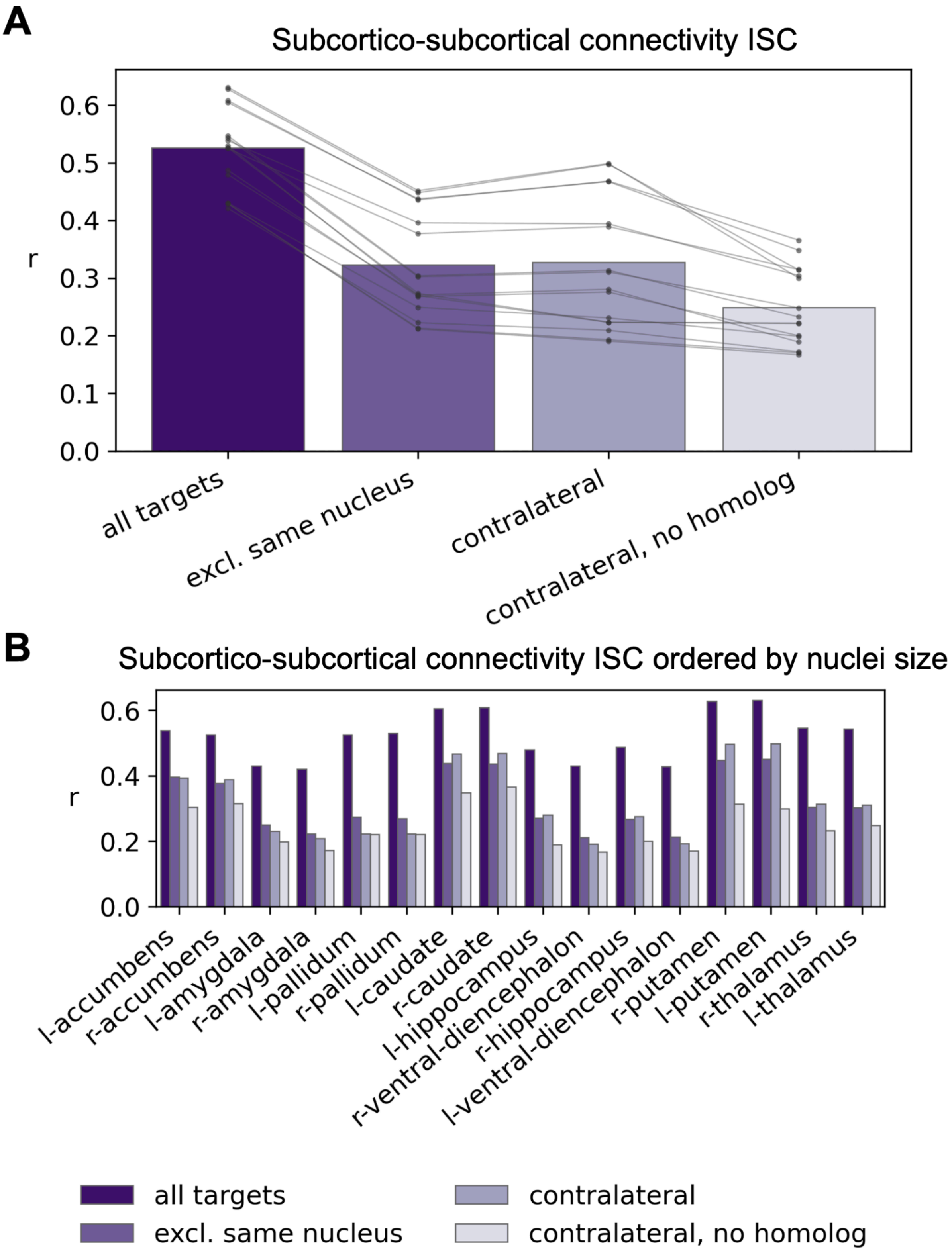


**Supplementary Figure 5.** *Ablation analysis of intersubject correlation (ISC) in subcortico-subcortical connectivity in the "Raiders" dataset (average across 13*

*subjects).* A) Summary across all nuclei (each nucleus is one data point), B) Same data but plotted separately for each nucleus. All targets = no exclusion. Excl. same nucleus = excluding all voxels within the same nucleus. Contralateral = excluding the same hemisphere. Contralateral, no homolog = excluding the same hemisphere, and also the homolog nucleus.

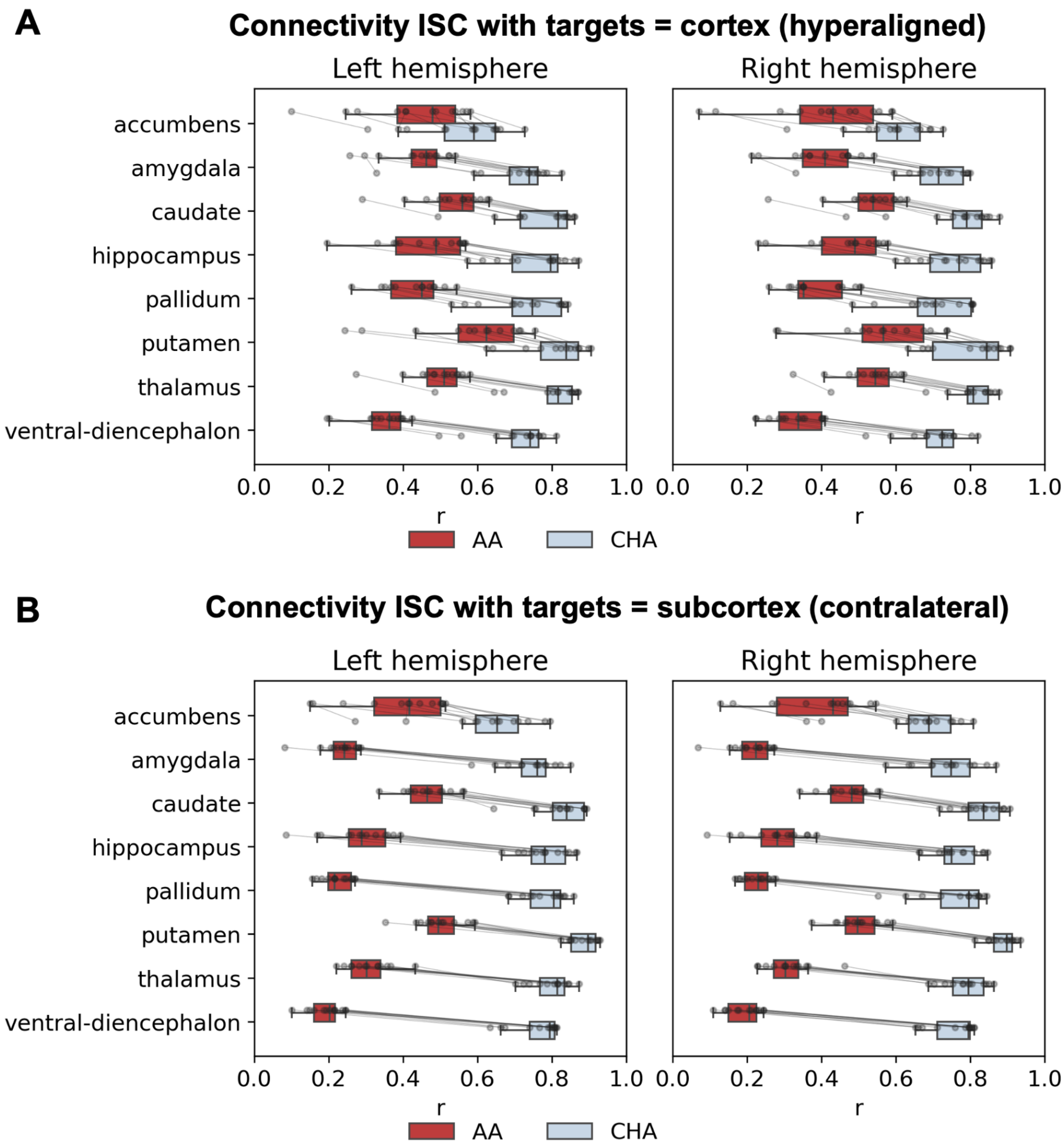


**Supplementary Figure 6.** *Leave-one-out connectivity intersubject correlation (ISC) for each subcortical nucleus in the Raiders dataset (13 subjects) with different targets:* A) hyperaligned cortical vertices (onavg-ico32); B) contralateral subcortical voxels. AA: Anatomical alignment only. CHA: Connectivity hyperalignment. Each data point represents an individual subject.

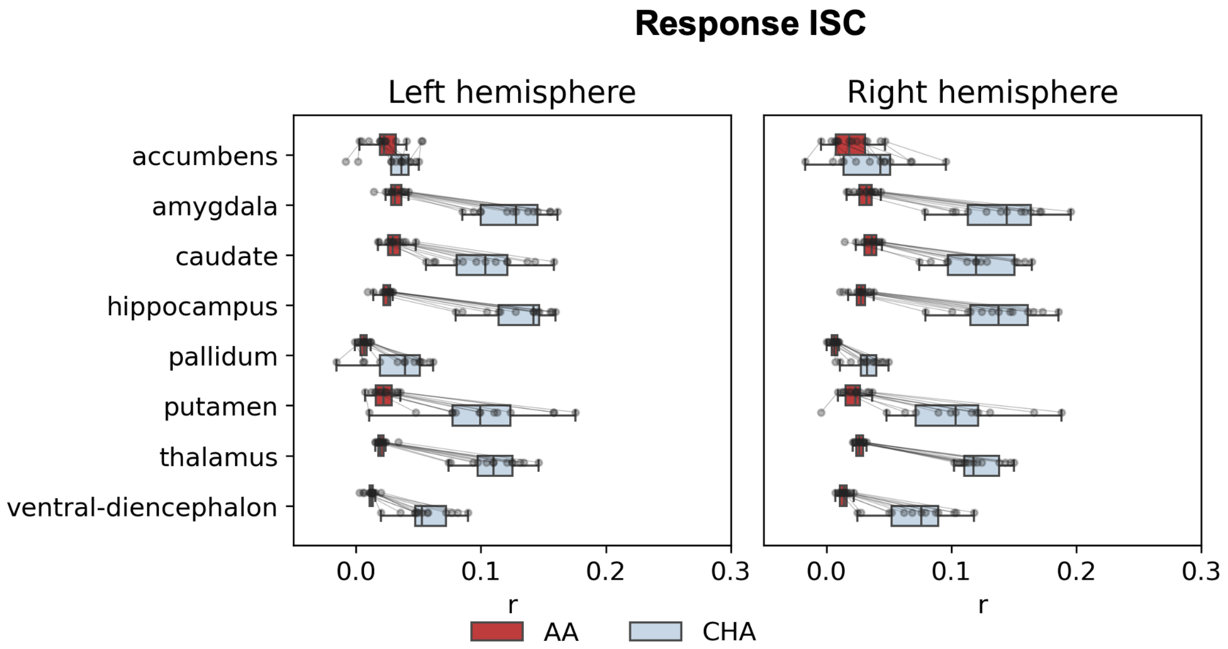


**Supplementary Figure 7.** *Leave-one-out response intersubject correlation (ISC) for each subcortical nucleus in the Raiders dataset (13 subjects).* AA: Anatomical alignment only. CHA: Connectivity hyperalignment. Each data point represents an individual subject.

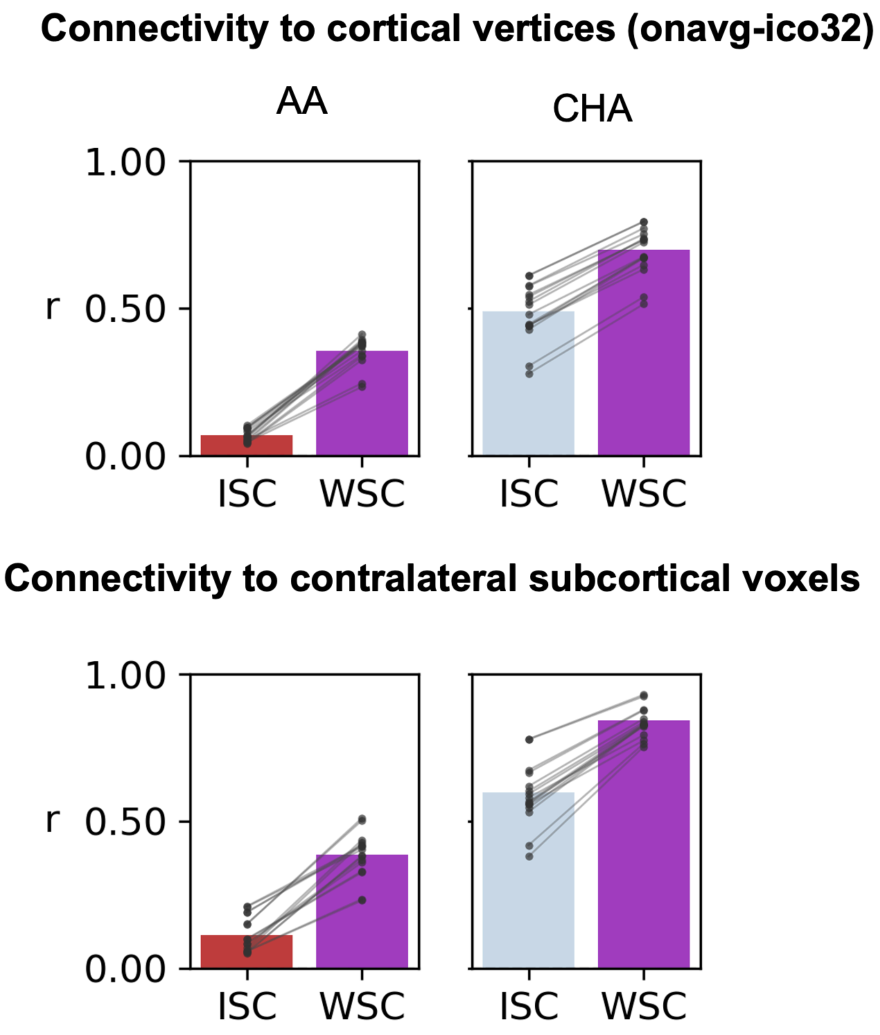


**Supplementary Figure 8.** *Within-subject correlation (WSC) and intersubject correlation (ISC) in connectivity across two runs in subcortical nuclei.* ISC was averaged across pairs for each subject and then averaged across subjects; WSC was averaged across subjects. AA: Anatomical alignment only. CHA: Connectivity hyperalignment. Each data point represents a subcortical nucleus.

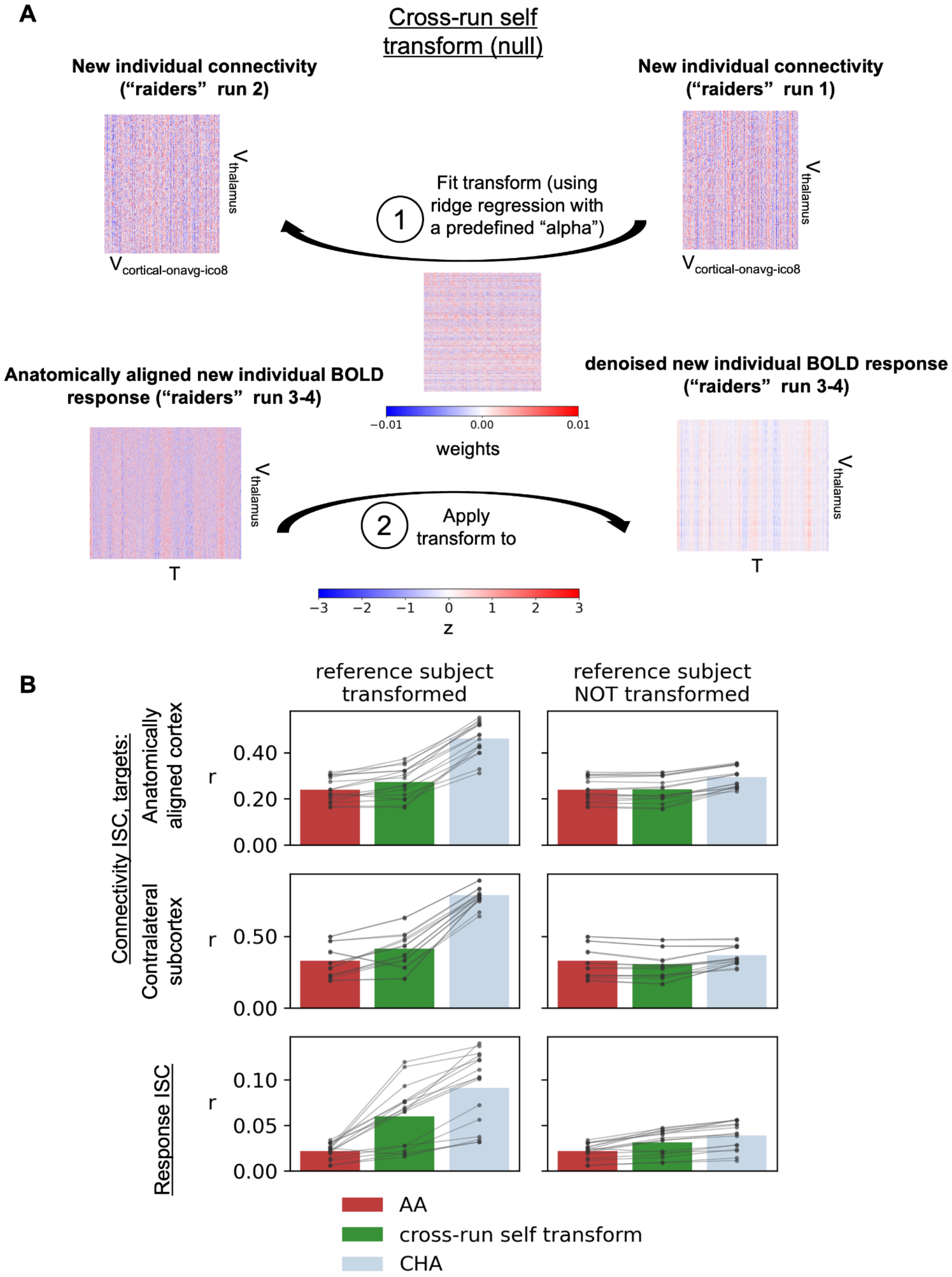
A
Cross-run self transform (null)
New individual connectivity ("raiders" run 2)
New individual connectivity ("raiders" run 1)
Fit transform (using ridge regression with a predefined "alpha")
1
V thalamus
V cortical-onavg-ico8
Anatomically aligned new individual BOLD response ("raiders" run 3-4)
denoised new individual BOLD response ("raiders" run 3-4)
−0.01 0.00 0.01
weights
2
Apply transform to
T
−3 −2 −1 0 1 2 3
z
B
reference subject transformed
reference subject NOT transformed
Connectivity ISC, targets:
Anatomically aligned cortex
Contralateral subcortex
Response ISC
r
0.00 0.20 0.40
0.00 0.50
0.00 0.05 0.10
AA
cross-run self transform
CHA

**Supplementary Figure 9.** *Hyperalignment to template or to subject with cross-run self-transform nulls.* A) Cross-run self-transform null was calculated by hyperaligning the run1 connectivity to run2 connectivity for the same subject with ridge regression, with a preselected alpha to match the actual hyperalignment. B) Reference subject transformed: leave-one-out intersubject correlation (looISC) where all subjects (including the reference subject) were applied a single transform to template (CHA) or cross-run self-transform with matched alpha. Reference subject NOT transformed: looISC where all but the reference subject were applied a single transform to the reference subject or cross-run self-transform with matched alpha. AA: anatomical alignment. CHA: connectivity hyperalignment.

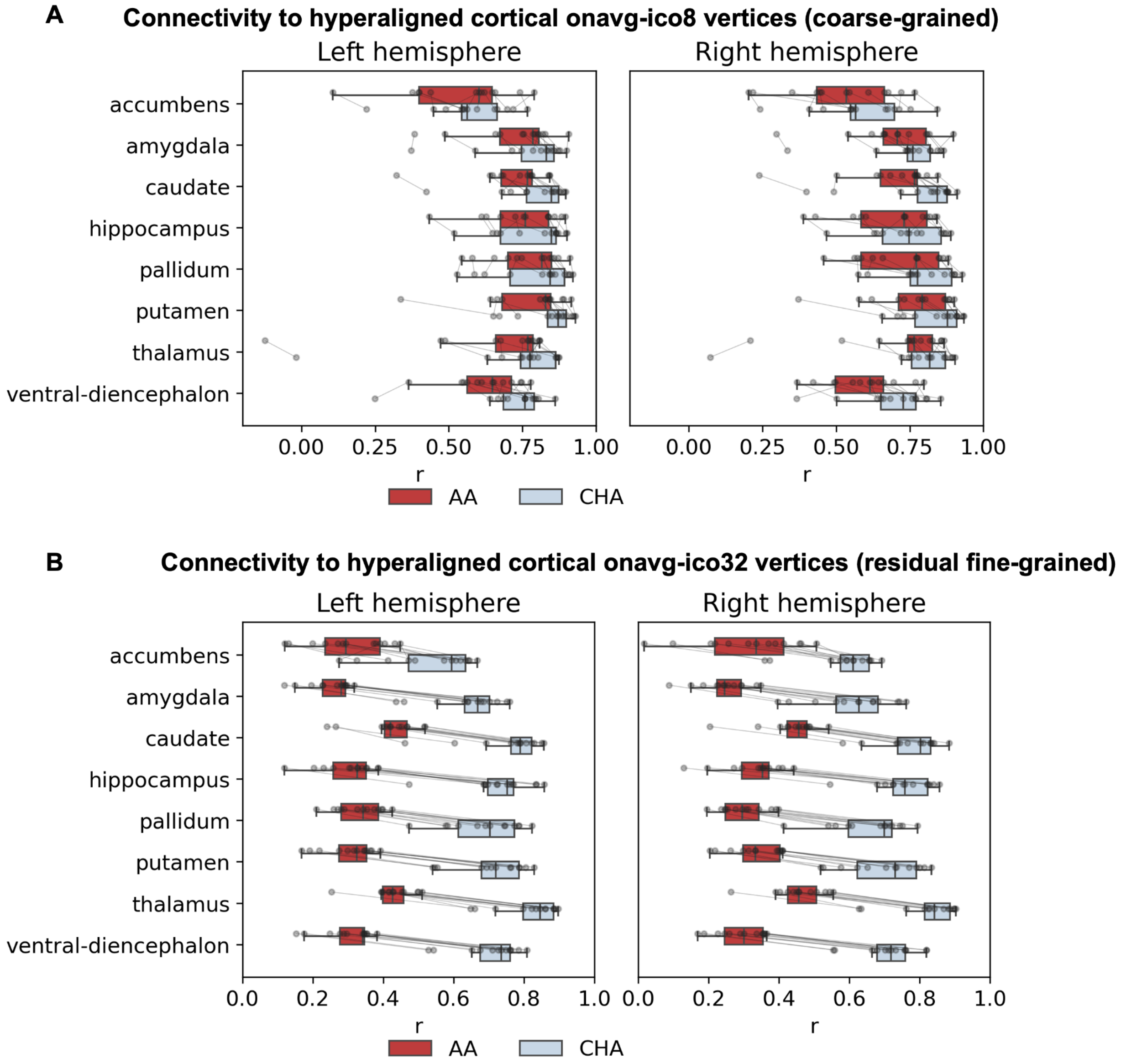

**Supplementary Figure 10**. *Leave-one-out connectivity intersubject correlation (ISC) for each subcortical nucleus in 13 subjects in A) coarse-grained and B) residual fine-grained connectivity to all hyperaligned cortical vertices (onavg-ico8 for coarse and onavg-ico32 for fine).* AA: Anatomical alignment only. CHA: Connectivity hyperalignment. Each data point represents an individual subject.

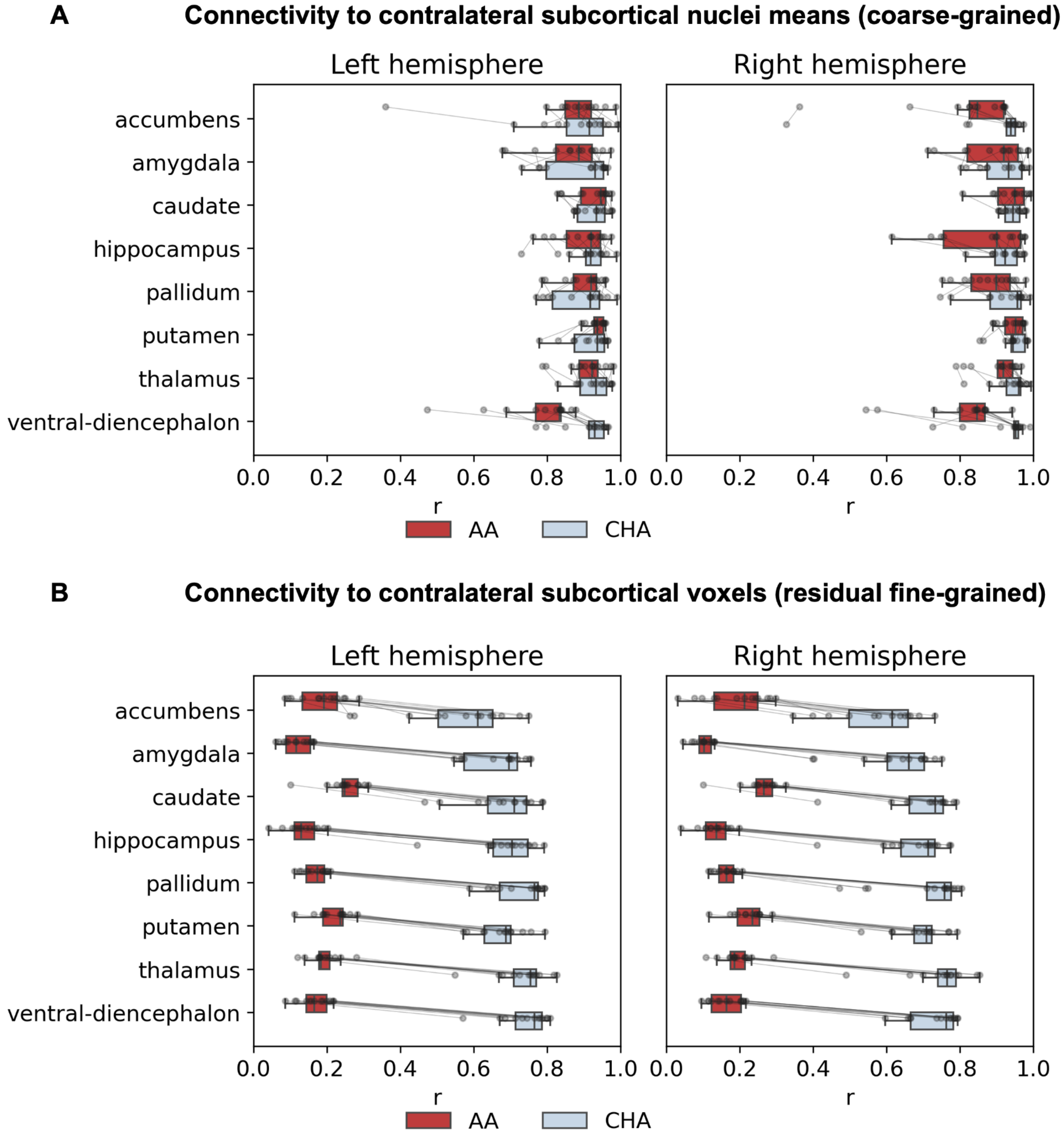


**Supplementary Figure 11**. *Leave-one-out connectivity intersubject correlation (ISC) for each subcortical nucleus in 13 subjects in A) coarse-grained connectivity to contralateral subcortical nuclei means and B) residual fine-grained connectivity to*

*contralateral subcortical voxels.* AA: Anatomical alignment only. CHA: Connectivity hyperalignment. Each data point represents an individual subject.

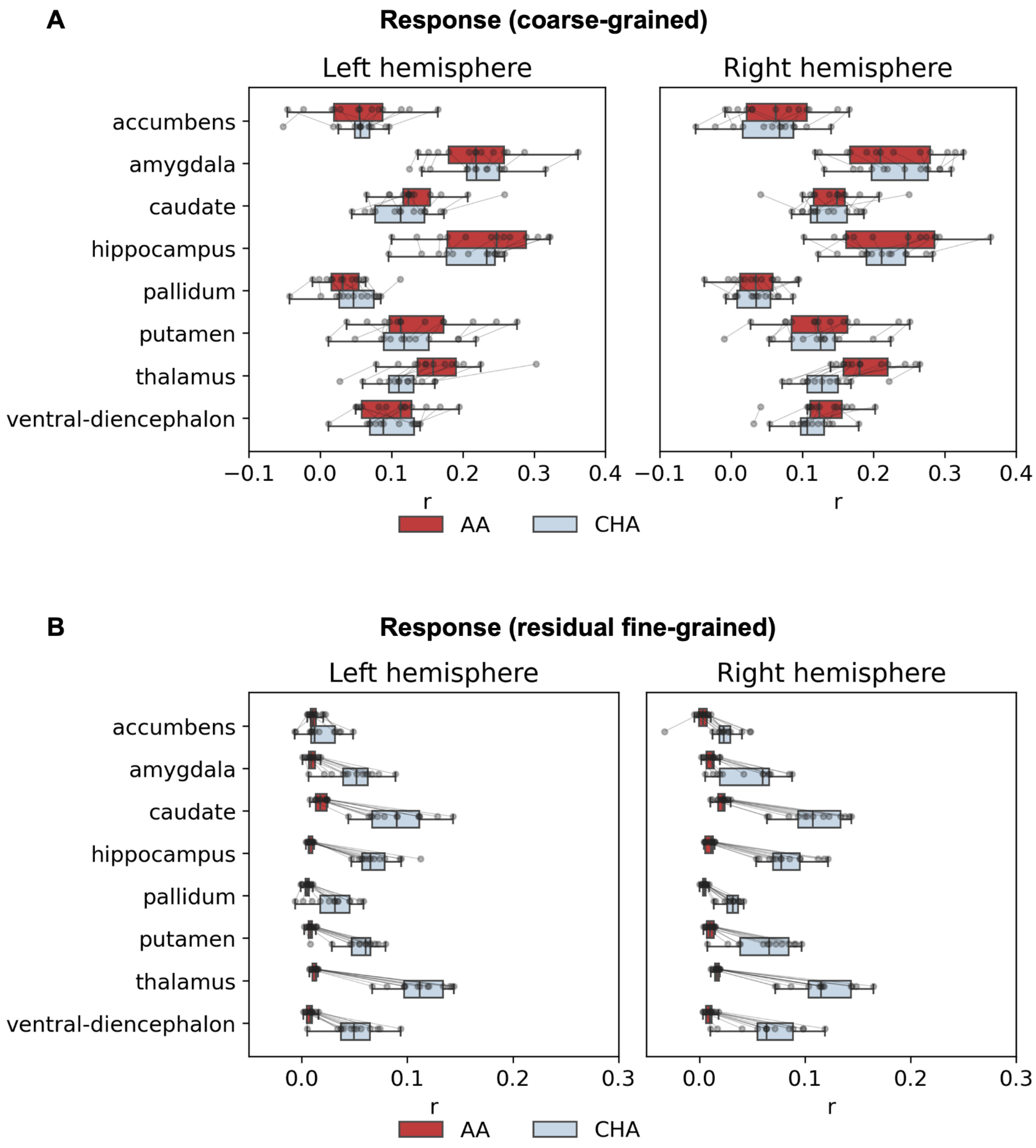


**Supplementary Figure 12**. *Intersubject correlation (ISC) for each subcortical nucleus in 13 subjects of* A) coarse-grained and B) residual fine-grained movie responses. AA: Anatomical alignment only. CHA: Connectivity hyperalignment. Each data point represents an individual subject.

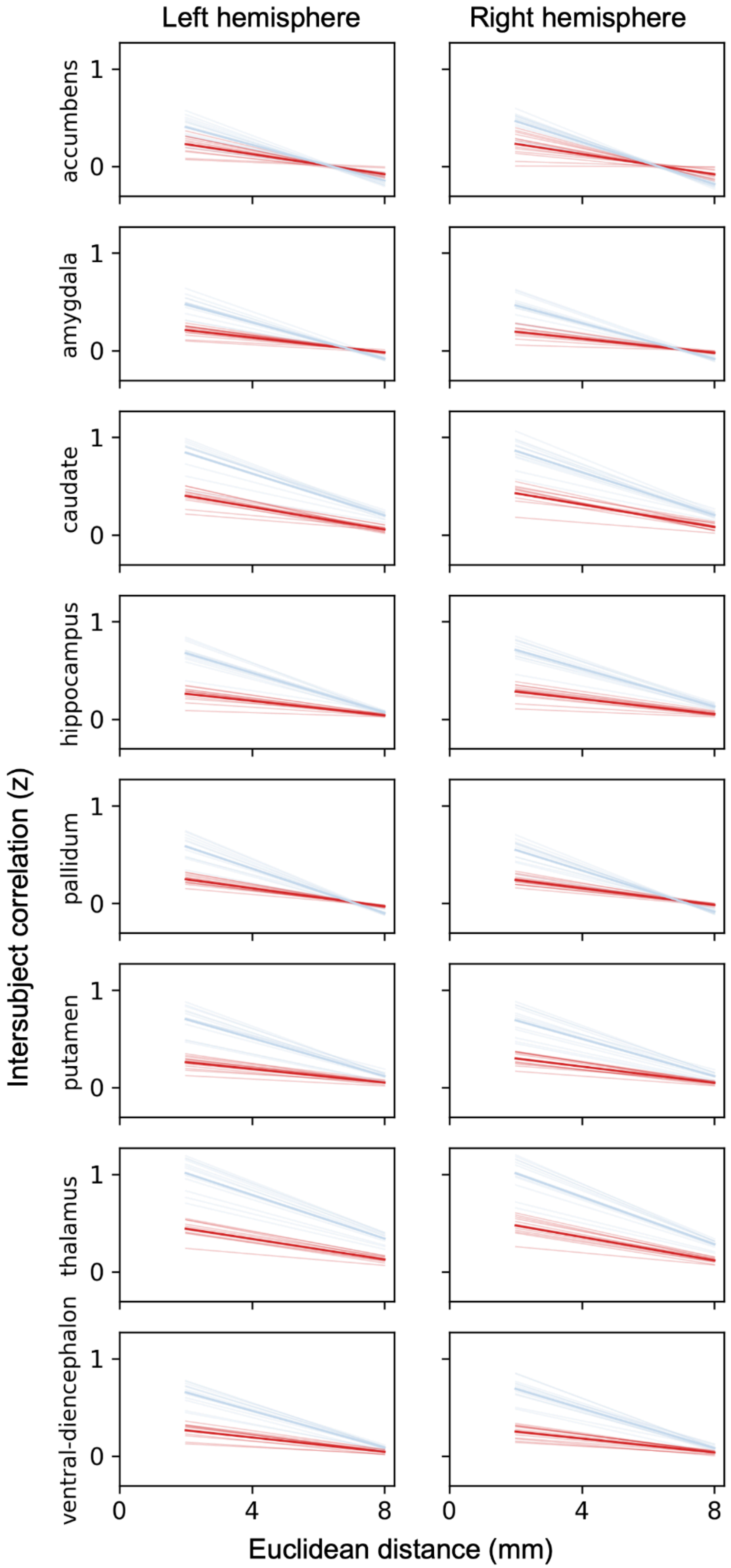
Left hemisphere
Right hemisphere
Intersubject correlation (z)
accumbens
amygdala
caudate
hippocampus
pallidum
putamen
thalamus
ventral-diencephalon
0
1
0
4
8
Euclidean distance (mm)

**Supplementary Figure 13**. *Intersubject spatial point spread function for ISC (z) of residual fine-grained connectivity to hyperaligned cortical (onavg-ico32) vertices.* Mean and standard error across 13 subjects. Each thin line is one subject, and the thick line is the average.

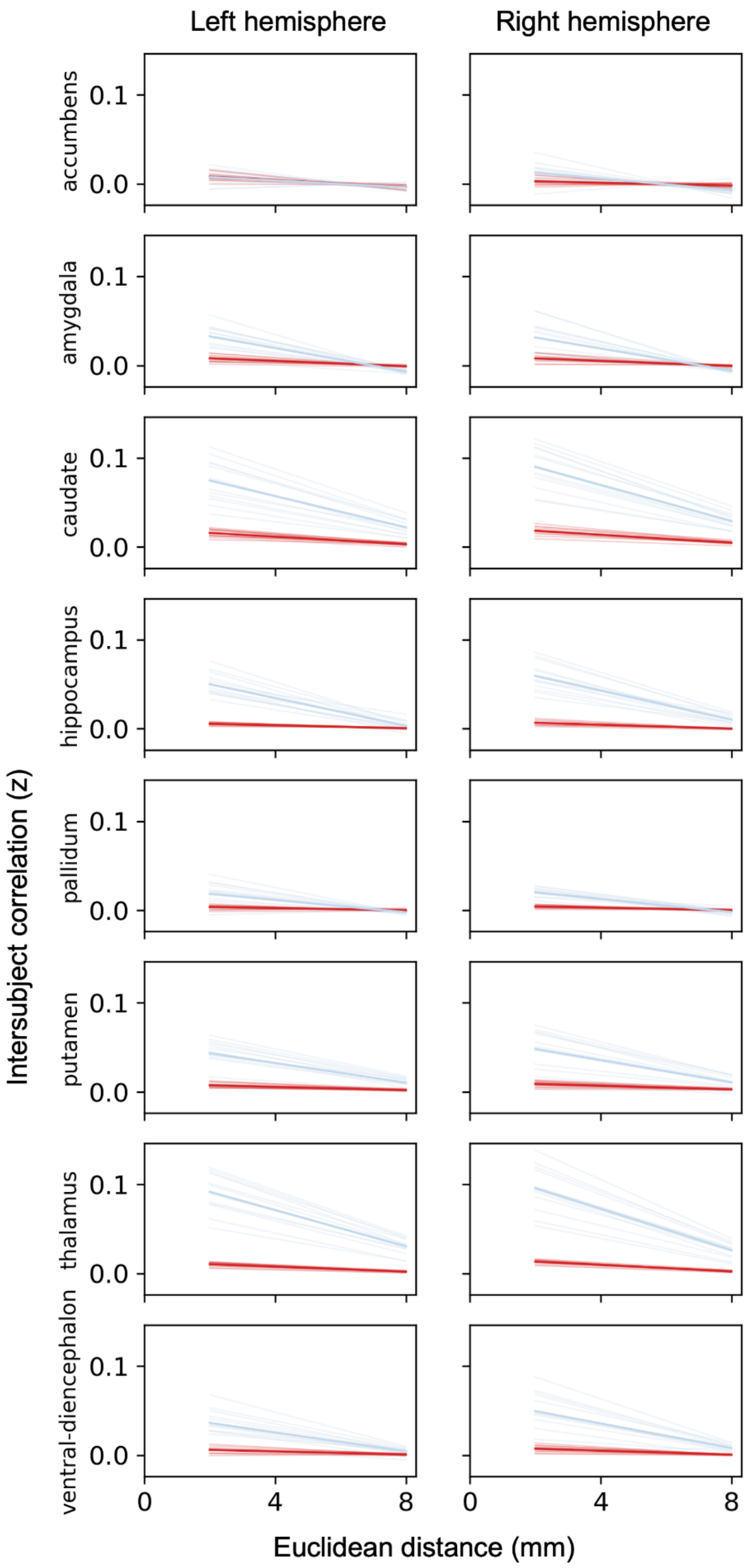

Left hemisphere
Right hemisphere
accumbens
amygdala
caudate
hippocampus
pallidum
putamen
thalamus
ventral-diencephalon
0.1
0.0
0
4
8
Intersubject correlation (z)
Euclidean distance (mm)

**Supplementary Figure 14**. *Intersubject spatial point spread function for ISC (z) of residual fine-grained connectivity to contralateral subcortical voxels.* Each thin line is one subject, and the thick line is the average.

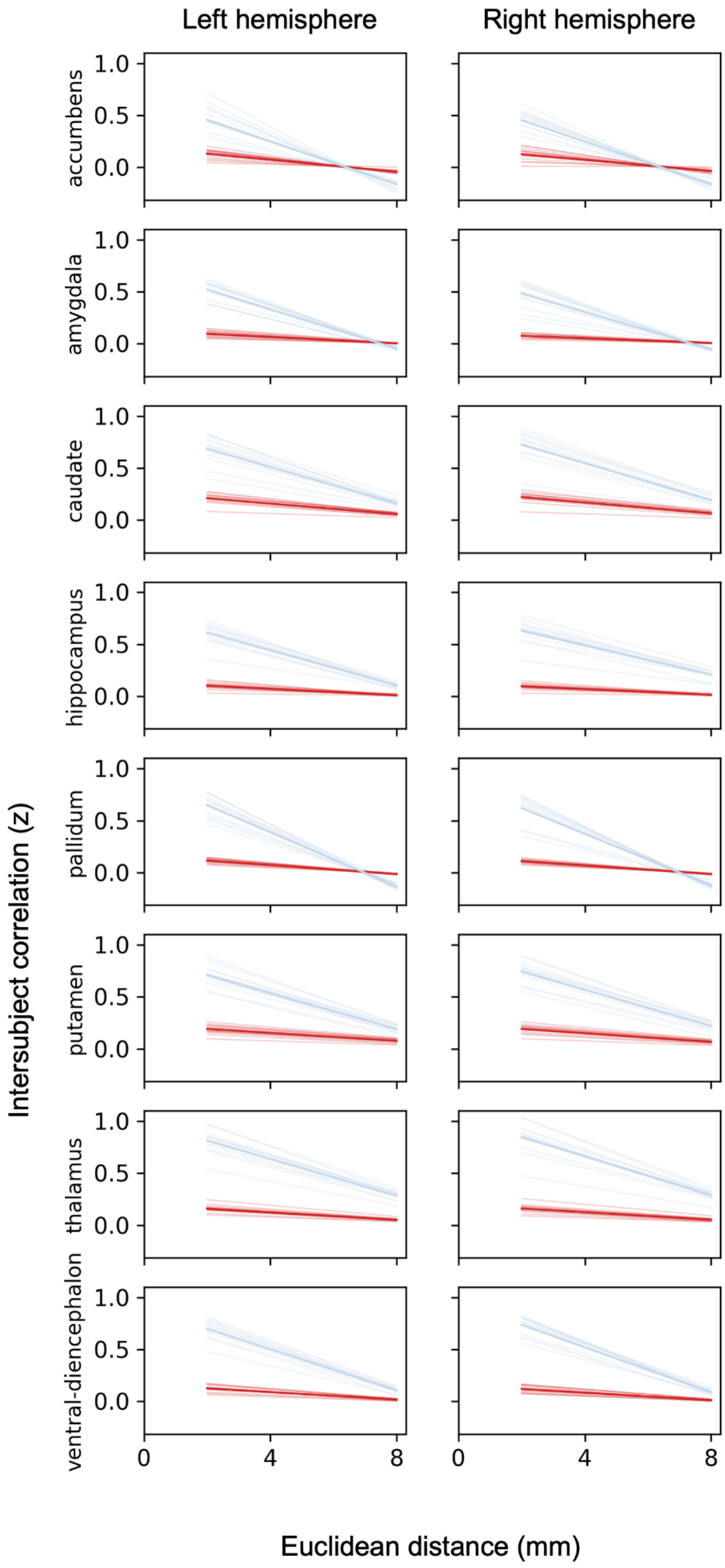

Left hemisphere
Right hemisphere
accumbens
amygdala
caudate
hippocampus
pallidum
putamen
thalamus
ventral-diencephalon
1.0
0.5
0.0
0
4
8
Intersubject correlation (z)
Euclidean distance (mm)

**Supplementary Figure 15**. *Intersubject spatial point spread function for ISC (z) of residual fine-grained responses.* Each thin line is one subject, and the thick line is the average.

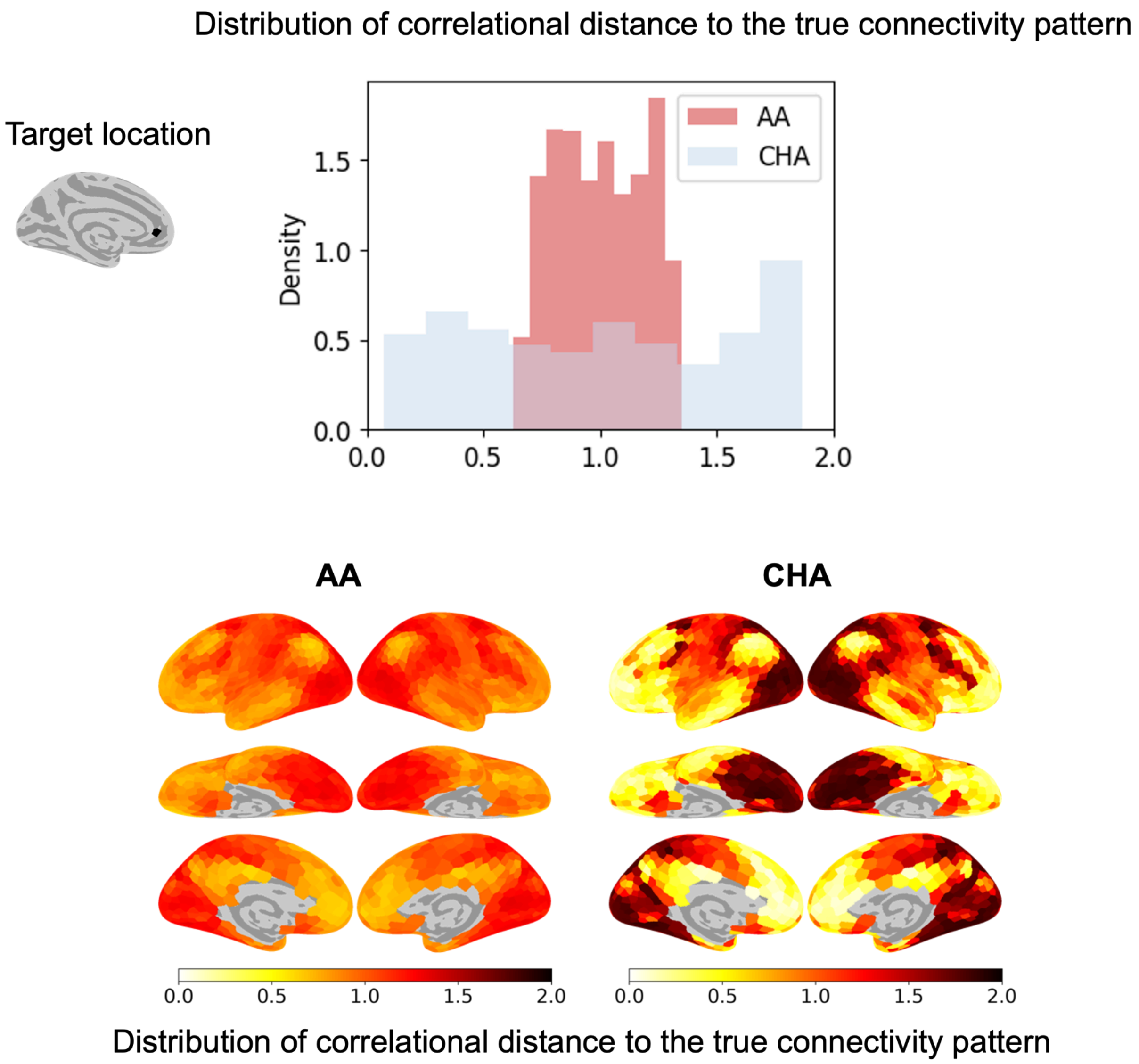


**Supplementary Figure 16.** *Example connectivity target (1210 onavg-ico8 cortical targets) prediction across subjects based on the residual fine-grained multivoxel pattern in each subcortical nucleus under anatomically aligned (AA) and connectivity-hyperaligned (CHA) conditions.* The histogram shows the distribution of correlational distance between the left thalamus connectivity pattern of the test subject at a given target location and the left thalamus connectivity patterns to each of the 1210 onavg-ico8 cortical targets in other subjects' average. The brain plots show the same correlational distance values visualized on brain surfaces. AA: anatomically aligned subcortical data and cortical data. CHA: connectivity hyperaligned subcortical data and cortical data.

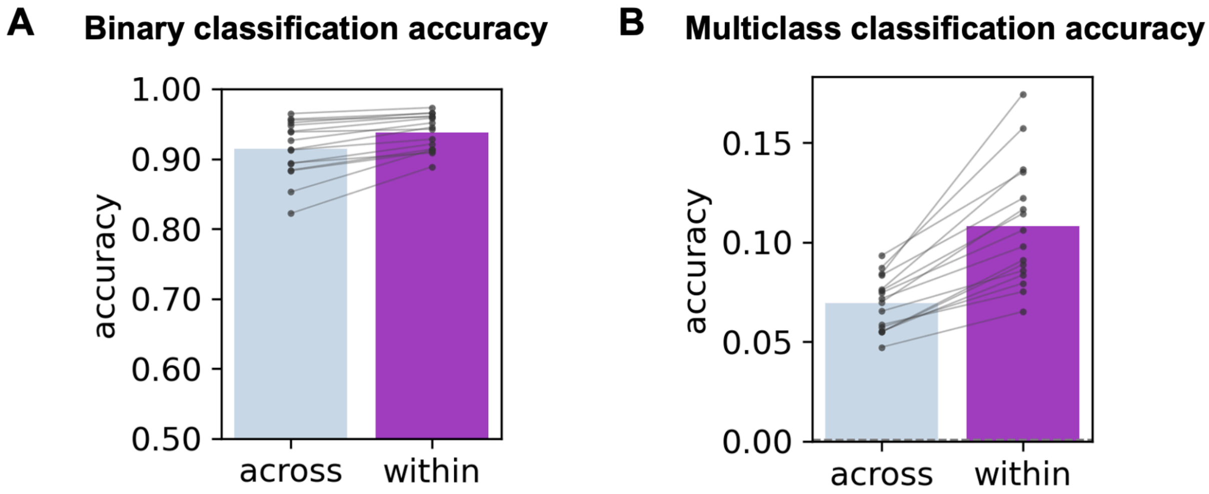


**Supplementary Figure 17.** *Connectivity target prediction across-subject versus within-subject (after connectivity hyperalignment).* A) Binary classification accuracy. B) Multiclass classification accuracy. Each dot and line represents one of the 16 nuclei.

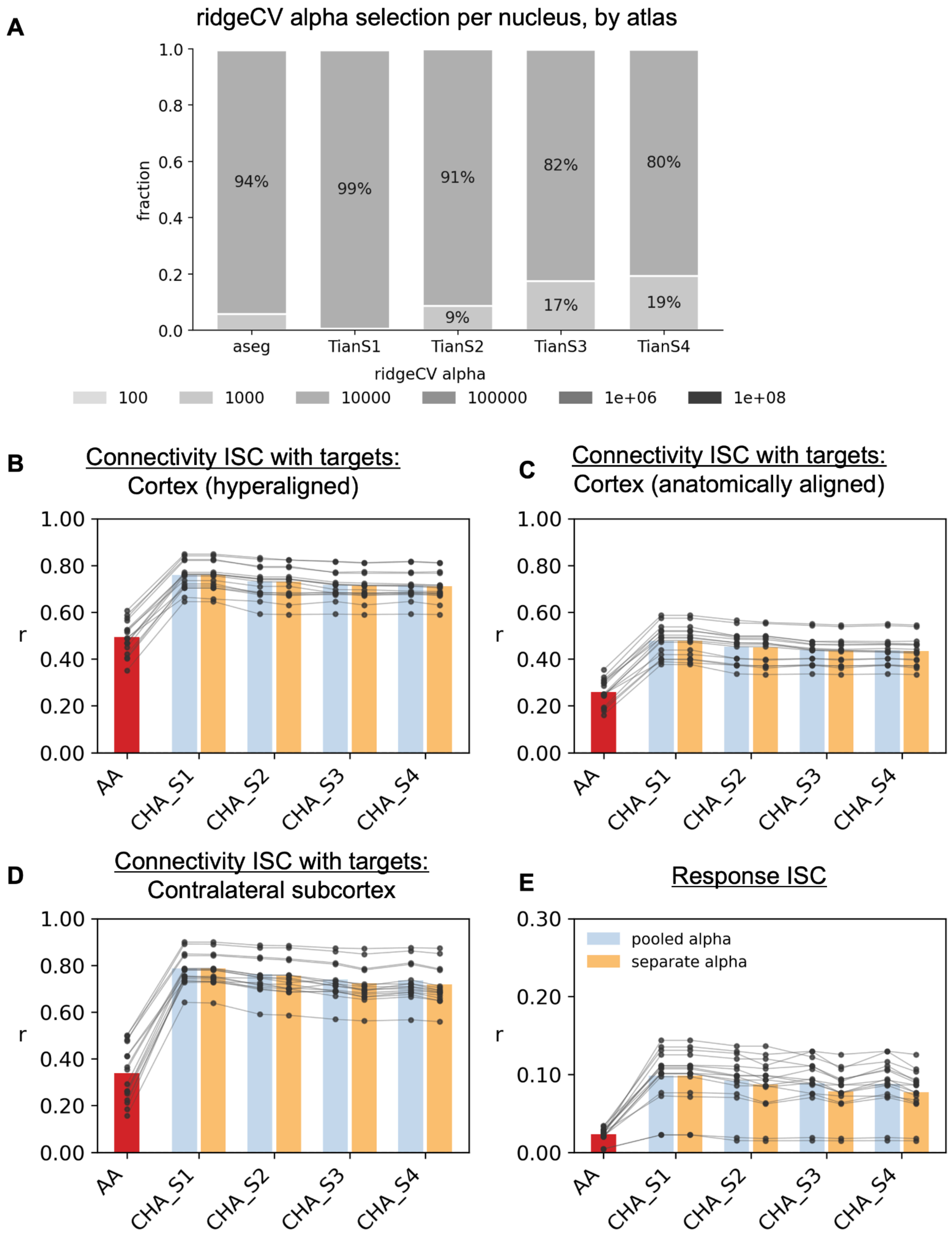


**Supplementary Figure 18.** *Replication of Figure 6 using per-nucleus (separate) ridge alpha selection compared to whole-brain (pooled) alpha selection.* A) The frequency distribution of alpha chosen by cross-validation across all subject-nucleus combinations. For comparison, pooled alpha was 10000 under all atlases. ISC in full fine-grained connectivity to B) hyperaligned cortical vertices (onavg-ico32), C) anatomically aligned cortical vertices (onavg-ico32), and D) contralateral subcortical voxels. E) ISC in full fine-grained movie responses. AA: anatomical alignment only. CHA: connectivity hyperalignment. Each data point represents one nucleus.
